# Loss of SUV420H2 promotes EGFR inhibitor resistance in NSCLC through upregulation of MET via LINC01510

**DOI:** 10.1101/2020.03.17.995951

**Authors:** A.S. Pal, A.M. Agredo, N.A. Lanman, J. Clingerman, K Gates, A.L. Kasinski

## Abstract

Epidermal growth factor receptor inhibitors (EGFRi) are standard-of-care treatments administered to patients with non-small cell lung cancer (NSCLC) that harbor EGFR alterations. However, development of resistance within a year post-treatment remains a major challenge. Multiple mechanisms can promote survival of EGFRi treated NSCLC cells, including secondary mutations in EGFR and activation of bypass tracks that circumvent the requirement for EGFR signaling. Nevertheless, mechanisms involved in bypass track activation are understudied, and in a subset of cases the mechanisms are unknown. The findings from this study identified an epigenetic factor, SUV420H2 that when lost drives resistance of NSCLC to multiple EGFRi, including erlotinib, gefitinib, afatinib, and osimertinib. SUV420H2 catalyzes trimethylation of histone H4 lysine-20, a modification required for gene repression and maintenance of heterochromatin. Here we show that loss of SUV420H2 leads to upregulation of an oncogenic long non-coding RNA, *LINC01510* that promotes transcription of the oncogene MET, a component of a major bypass track involved in EGFRi resistance.

**Significance:** Due to an incomplete understanding of the mechanisms involved in promoting resistance to EGFRi, patients often succumb to their disease. Here we identified a global mediator of EGFRi resistance, SUV420H2 that helps to uncover an additional mechanism involved in resistance driven via a major bypass track involving the protooncogene MET.

## Introduction

Lung cancer is the leading cause of cancer-related mortality, with an estimated 135,720 deaths predicted in 2020 in the United States alone (Siegel et al., 2020). The majority of lung cancer patients are diagnosed with non-small cell lung cancer (NSCLC), a subtype that represents 85% of lung cancer cases. Since most lung cancer patients are diagnosed at later stages with metastatic disease surgical resection is not curative, and thus, the most effective treatment strategies are radiotherapy, chemotherapy, and targeted therapy. Targeted therapeutics are selected based on the presence of particular molecular drivers, genes that the cancer cells are essentially addicted to. A few such drivers that are commonly present in NSCLC include KRAS, MEK, MET, HER2, and EGFR, many of which are either mutated or amplified, resulting in constitutive pro-growth signaling (Luo et al., 2008; Onitsuka et al., 2010).

Epidermal Growth Factor Receptor (EGFR) is a cell surface receptor required for normal cell growth and proliferation. In 10-35% of NSCLC cases EGFR and its downstream pro-growth signaling pathways are constitutively activated due to mutations in the receptor, the most common of which are an amino acid substitution in exon 21 (L858R) or an in-frame deletion in exon 19. Mutant EGFR can be clinically targeted with a variety of EGFR tyrosine kinase inhibitors (EGFRi), including erlotinib and gefitinib, both of which are first generation EGFRi, afatinib, a second-generation inhibitor, or osimertinib a recently approved third-generation EGFRi. Osimertinib not only targets the primary EGFR mutation but is also active against a secondary mutation in EGFR, T790M. Currently, EGFRi are front-line therapies for patients with tumors that have the respective EGFR mutations, but erlotinib has also been approved for use in patients without EGFR mutations that have failed at least one prior chemotherapy regimen (Osarogiagbon et al., 2015). Erlotinib binds reversibly and specifically to the ATP-binding pocket of EGFR with high efficacy, abrogating downstream growth and survival signaling pathways. While initially beneficial as a first- or second-line treatment, patients develop resistance within a year post treatment, which is currently a major drawback to its use (Liao et al., 2019). Indeed, a similar response was observed with imatinib, another targeted inhibitor used to treat patients with chronic lymphocytic leukemia (Gambacorti-Passerini et al., 2003). The targeted gene either incurs additional mutations or activates alternative signaling pathways to evade therapy. In the case of erlotinib treated patients that develop resistance, over 60% of patients acquire a secondary mutation, T790M, in their tumor whereas approximately 20% of tumors utilize bypass tracks. Bypass tracks allow the tumor to escape inhibition of the EGFR pathway through the use of alternative mechanisms that sustain their survival. Bypass tracks include signaling through oncogenic proteins such as MET, BRAF, HER2, PIK3CA or histological transformation of cells - NSCLC into small cell lung cancer (SCLC) or through epithelial to mesenchymal transition (EMT) (Liao et al., 2019; Niederst et al., 2015; Sequist et al., 2011; Yun et al., 2008). In addition to an incomplete understanding of mechanisms that govern these bypass tracks there are also approximately 15-20% of NSCLC tumors that acquire erlotinib resistance by mechanisms that remain unidentified (Liao et al., 2019).

While gain-of-function mechanisms that drive resistance have been identified, loss of tumor suppressive genes, such as PTEN, TP53, TET1, and NF1 has also been reported to contribute to resistance (de Bruin et al., 2014; Forloni et al., 2016; Huang et al., 2011; Sos et al., 2009). Indeed, many tumor suppressive proteins function as gatekeepers of the genome preventing spurious activation of oncogenes. To better define genes that prevent the development of resistance, a genome-wide loss of function screen was conducted using the CRISPR-Cas9 system. Our data suggest that an epigenetic factor and *bona fide* tumor suppressor, SUV420H2 can be included among the gatekeepers of the genome. SUV420H2 catalyzes the trimethylation of histone H4 at lysine-20 (H4K20) using mono-methylated H4K20 as a substrate, which is required for the establishment of heterochromatin and repression of genes (Hahn et al., 2013; Schotta et al., 2008; Weirich et al., 2016). Loss of SUV420H2 has previously been implicated in causation of multiple cancers (Fraga et al., 2005; Pogribny et al., 2007), but for the first time we show that loss of SUV420H2 is a novel mechanism that promotes erlotinib resistance in NSCLC cells. The findings of this study determined that SUV420H2 mutant cells express high levels of the oncogenic long non-coding RNA, *LINC01510* that transcriptionally upregulates the oncogene MET, mediating erlotinib resistance.

## Results

### Identification of mediators of erlotinib resistance

To identify genes, that when mutated confer resistance to erlotinib sensitive cells, a genome-wide CRISPR-Cas9 knock-out screen was performed. The screen was conducted in EKVX cells, a cell line that was determined to be erlotinib sensitive in data obtained from the Developmental Therapeutics Program, which is maintained by the National Cancer Institute (NCI-60, DTP (NCI-60 DTP)). The cells were engineered to stably express the Cas9 protein and resulting clones were validated for their response to erlotinib, which was similar to the parental EKVX cells **(Supplementary Figure 1).** A single Cas9-expressing EKVX clone was taken forward to conduct the screen, which is hereafter referred to as ECas9. The ECas9 cells were infected with the GeCKO V2 sgRNA lentiviral library targeting 19,052 protein-coding genes and 1,864 miRNA genes (**Figure 1A**) (Shalem et al., 2015). To obtain full coverage of the lentiviral sgRNA library, transduction was performed at 300-fold coverage and was conducted in triplicates to mitigate false positives. One third of the transduced cells were used to determine the representation of the integrated sgRNAs prior to selection (baseline). The remaining cells were grown for 15 passages in the presence of 1.23 uM erlotinib, a concentration that inhibits the growth of 75% of cells (GI75). Integrated sgRNAs were identified from the resulting population and from the baseline cells by PCR amplification and subsequent high-throughput sequencing. Combined analysis of the three replicates using the MAGeCK-VISPR algorithm identified significantly enriched sgRNAs in the population of cells that were cultured in erlotinib (**Supplementary Table 1**, **Figure 1B**) (Liu et al., 2014). Following the analysis, multiple genes that were previously reported to be 1) downregulated during acquired resistance to chemotherapy treatment (EGFRi or non-EGFRi) (Chen et al., 2016), 2) highly expressed in erlotinib sensitive cells (CtRP v2; Orzáez et al., 2012), and 3) *bona fide* tumor suppressors (Aprelikova et al., 2013; Chen et al., 2011; Fraga et al., 2005; Kantidakis et al., 2016; Ryu et al., 2013; Shinchi et al., 2015), were identified among the top hits, validating the sensitivity of the screen.

**Figure 1:**
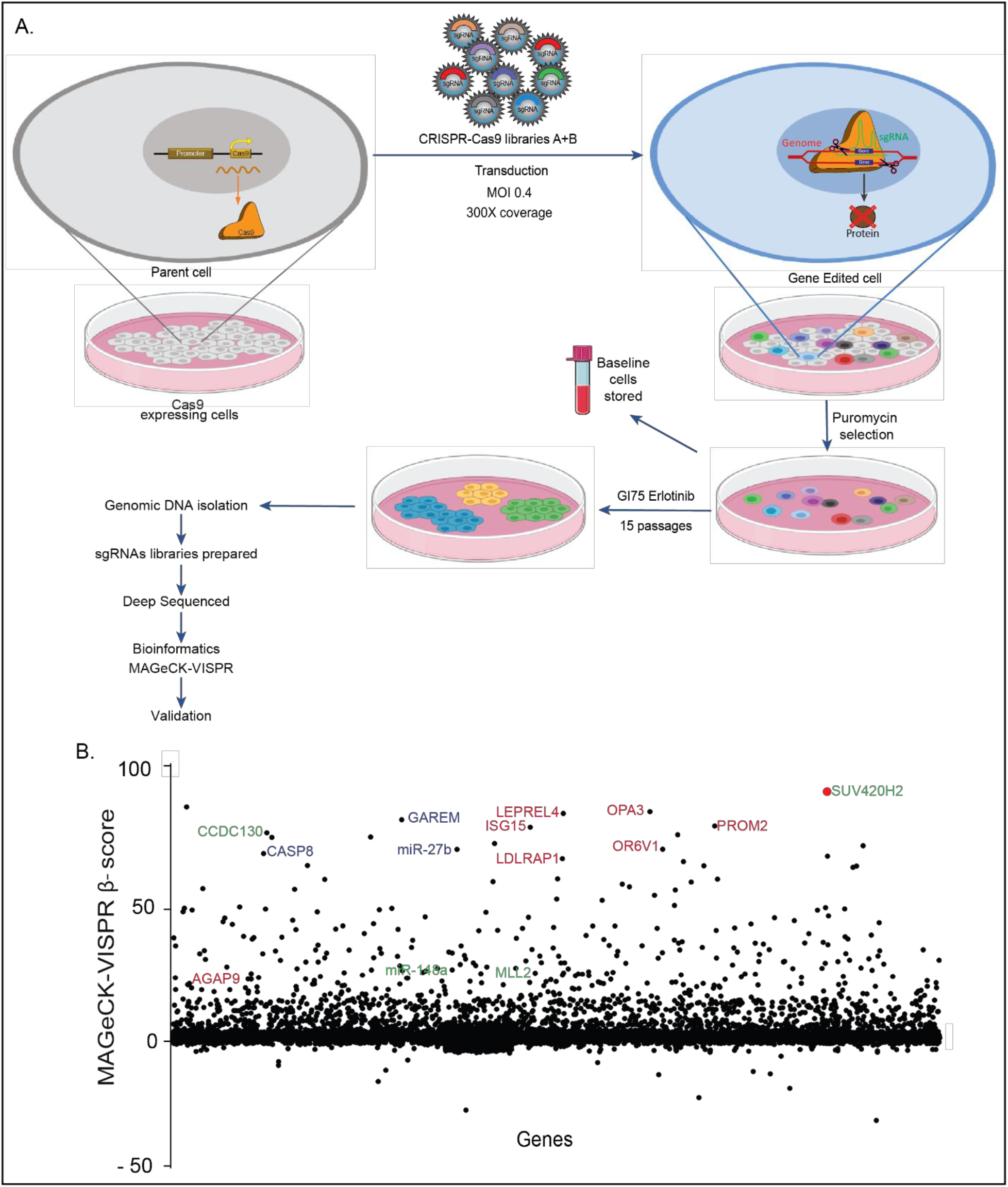
A genome-wide CRISPR-Cas9 screen identifies mediators of erlotinib resistance in NSCLC. A) Outline of the CRISPR-Cas9 knock-out screen. B) Fold enrichment (β-score) analysis of sgRNAs of the CRISPR-Cas9 lentiviral library targeting genes, evaluated using MAGeCK-VISPR analysis. Genes represented in blue have previously been reported to be downregulated in cells post chemotherapeutic treatment, in red are genes reported to be high in erlotinib sensitive cells (CtRPv2), and in green are previously reported tumor suppressors.

### Low expression of SUV420H2 is associated with erlotinib resistance, and predicts poor prognosis in NSCLC

The top hit from the CRISPR-Cas9 knock-out screen, SUV420H2 is a histone methyltransferase encoded by *KMT5C*. SUV420H2 specifically trimethylates histone H4 lysine-20 (H4K20), which is associated with transcriptional repression and is important for establishing constitutive heterochromatic regions (Hahn et al., 2013; Schotta et al., 2008). Multiple studies have reported on the role of SUV420H2 as a tumor suppressor, and both SUV420H2 and H4K20 trimethylation (H4K20me3) are severely downregulated in multiple cancers (Van Den Broeck et al., 2008; Chekhun et al., 2007; Fraga et al., 2005; Pogribny et al., 2006; Shinchi et al., 2015). To determine if SUV420H2 is also a mediator of erlotinib response, various validation assays were performed. Firstly, using a panel of NSCLC cell lines, a negative correlation between *KMT5C* and erlotinib response was determined (**Figure 2A-C**, Pearson r = −0.83). Due to the lack of a sensitive and specific SUV420H2 antibody, the downstream effector of SUV420H2, H4K20me3 was evaluated as a proxy for SUV420H2 activity (**Supplementary Figure 2A,B)**. Indeed, in the same cell line panel, H4K20me3 levels positively correlate with *KMT5C* (Pearson r = 0.6905, **Supplementary Figure 2C**). Additionally, similar to the negative correlation between *KMT5C* and erlotinib response in the NSCLC panel, H4K20me3 also displayed a negative correlation with erlotinib response (Pearson r = −0.61, **Supplementary Figure 2D**). These strong correlations suggest a possible role for SUV420H2 and H4K20me3 levels in mediating the response of NSCLC cells to erlotinib.

**Figure 2:**
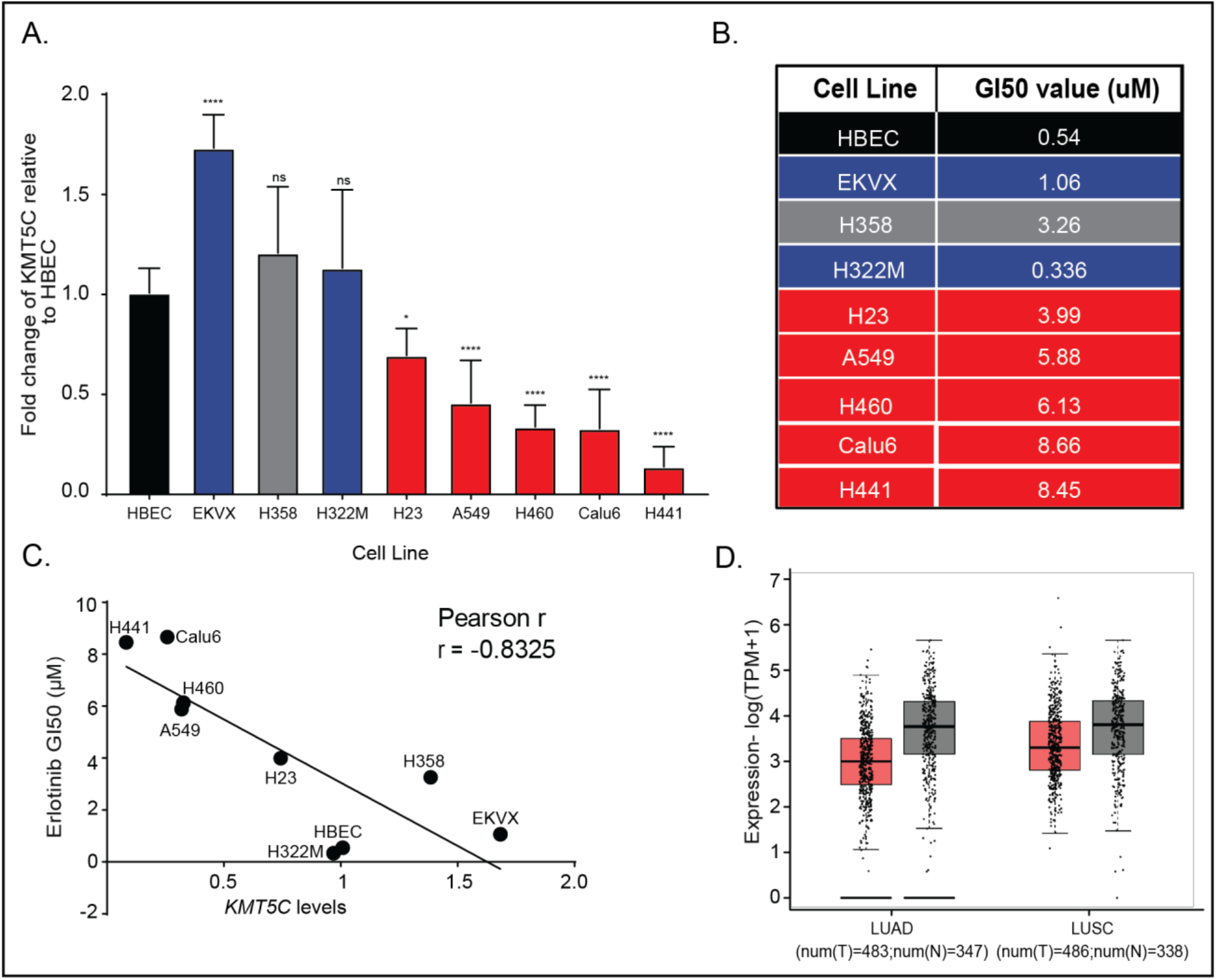
Reduced SUV420H2 expression correlates with erlotinib resistance in NSCLC cells, and poor prognosis in NSCLC patients. A) Expression of *KMT5C* in a panel of NSCLC cells, relative to a non-tumorigenic lung epithelial cell line (Human Bronchial Epithelial Cells, HBEC), evaluated by qRT-PCR. Data are normalized to GAPDH and are shown relative to levels in HBECs. One-way ANOVA followed by Dunnett’s Multiple Comparison test was used to evaluate statistical significance of *KMT5C* levels relative to HBECs. Color of bars represents previously reported response to erlotinib in the DTP study. Red bars indicated cells lines found to be resistant to erlotinib, blue bars represent cell lines reported to be sensitive. Neither HBEC (non-tumorigenic control) or H358 (NSCLC) were included in the DTP dataset. B) Erlotinib dose response was evaluated by exposing each of the individual cell lines to varying concentrations of erlotinib or the highest equivalent volume of DMSO (negative control) containing media for 72 hours followed by SRB assay. GI50 concentrations of erlotinib were calculated from each cell lines respective dose curve. Colors are as in A. C) Correlation analysis between *KMT5C* from A and GI50 erlotinib concentrations from B was evaluated using Pearson correlation test. D) GEPIA analysis for *KMT5C* levels in normal and tumor samples from LUAD and LUSC data obtained from The Cancer Genome Atlas (TCGA) and the Genotype-Tissue Expression (GTEx) databases. TPM= Transcripts per million, T= Tumor, N=Normal.

Next, we investigated *KMT5C* levels in NSCLC patient samples using publicly available data provided by The Cancer Genome Atlas (TCGA) and the Genotype-Tissue Expression (GTEx) projects. Patient samples were compared to non-cancerous control tissues using Gene Expression Profiling Interactive Analysis (GEPIA, **Figure 2D**) (Tang et al., 2017). *KMT5C* levels were generally lower in both lung adenocarcinoma (LUAD) and lung squamous cell carcinoma (LUSC) samples relative to normal samples suggesting SUV420H2 functions as a *bona fide* tumor suppressor.

### Loss of SUV420H2 confers resistance to EGFR inhibitors

To further validate the findings from the CRISPR-Cas9 screen, ECas9 cells (SUV420H2 wildtype) were transfected with a sgRNA targeting *KMT5C* to generate three SUV420H2 mutant lines, clones A, C and E. Genotyping validated that the sgRNA specifically targeted *KMT5C* resulting in various insertions and deletions (**Supplementary Figure 3A**). *KMT5C* levels were reduced in all three clones (**Figure 3A**) resulting in downregulation of H4K20me3 (**Figure 3B,3C** and **Supplementary Figure 3B**). Comparing the erlotinib response of the mutant clones to that of SUV420H2 wildtype cells confirmed that loss of SUV420H2 leads to 5.4 – 11.7 fold increase in erlotinib resistance (**Figure 3D**). Additionally, increased proliferation of the mutant clones relative to SUV420H2 wildtype cells in the presence of erlotinib corroborated the results (**Figure 3E**). We also evaluated the response of SUV420H2 mutant clones to other EGFRi including afatinib, gefitinib, and osimertinib. All the clones were resistant to all three EGFRi (**Supplementary Figures 3C-3H**) – resistance to the irreversible inhibitor, afatinib was ~300 to 900-fold higher in the mutant clones relative to wildtype SUV420H2 cells. Conversely, mutant clones were unaffected in the presence of cisplatin (data not shown) suggesting that loss of SUV420H2 is not a global mediator of resistance, but may be specific to EGFRi or perhaps other targeted agents.

**Figure 3:**
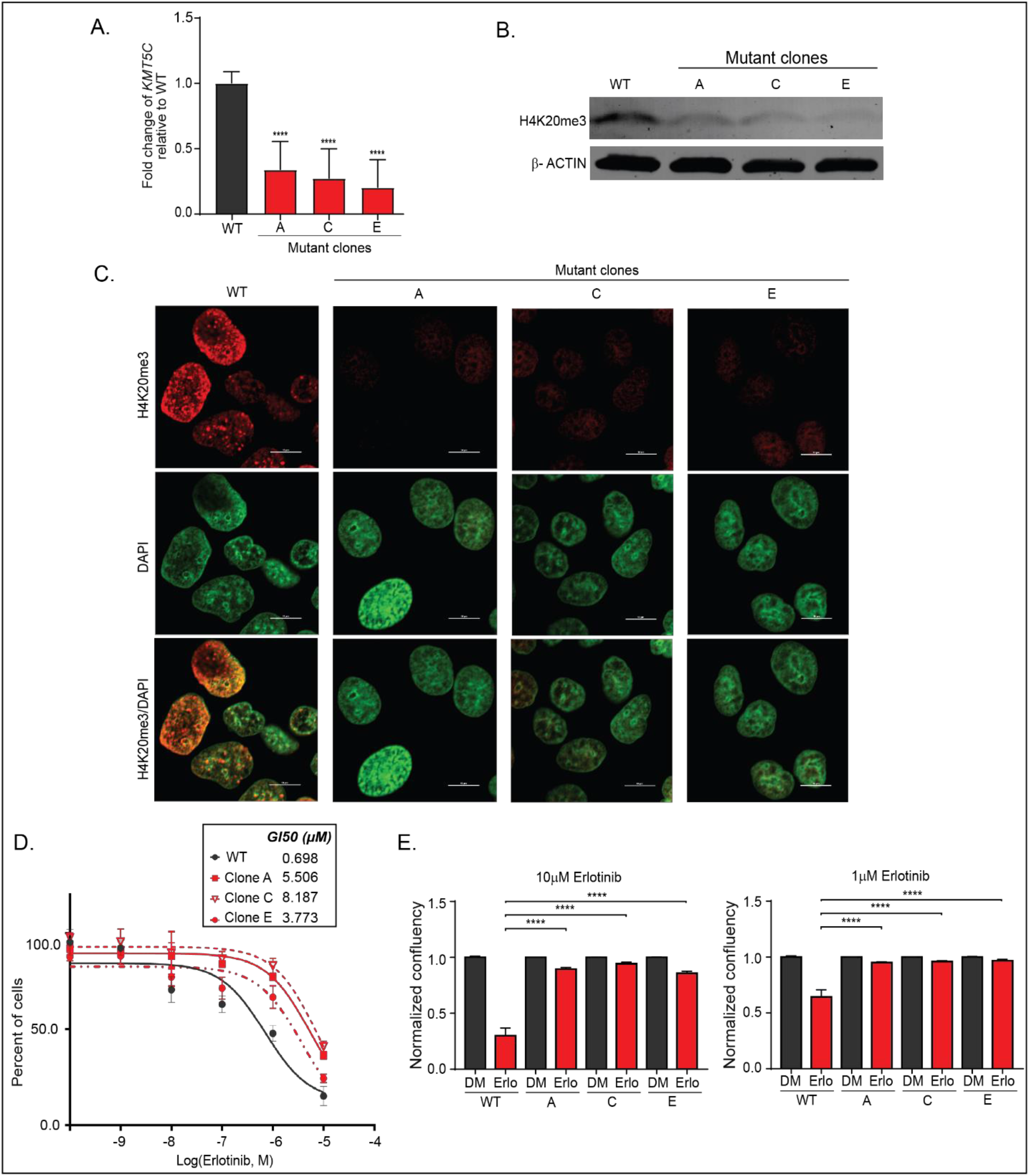
Loss of SUV420H2 confers resistance to erlotinib. A) Expression of *KMT5C* in mutant lines, clones A, C, E evaluated by qRT-PCR. Data are normalized to *GAPDH* and are represented relative to ECas9 (*KMT5C* wildtype, WT) cells. One-way ANOVA was used to evaluate statistical significance of *KMT5C* transcript levels relative to WT. B) Representative western blot of H4K20me3 in WT cells and mutant clones A, C, E. β-ACTIN serves as a loading control. C) Representative immunofluorescent image of H4K20me3 in WT cells and clones A, C, E. Scale bar, 10μm. D) Erlotinib dose response assayed using the SRB assay after exposing the indicated cells to varying concentrations of erlotinib or the highest equivalent volume of DMSO (negative control) containing media for 72 hours. Following normalization, the GI50 concentration of erlotinib was calculated from the respective dose curve for each cell line. E) Live cell imaging of WT or mutant clones A, C, E (represented as A, C, E) was conducted to quantify proliferating cells in the presence of erlotinib (Erlo) or vehicle control (DMSO, DM) for 72 hours. Data relative to respective normalized DMSO control treatments is represented. One-way ANOVA followed by Dunnett’s Multiple Comparison test was used to evaluate statistical significance of clones A, C, E in the presence of 10 or 1μM erlotinib compared to WT cells.

### Ectopic expression of SUV420H2 partially sensitizes EGFRi resistant cells

Since loss of SUV420H2 led to erlotinib resistance, we evaluated if the converse holds true by overexpressing SUV420H2. A doxycycline (DOX) inducible *KMT5C* plasmid was stably expressed in Calu6 cells, which have endogenously low levels of *KMT5C* (**Figure 2A**) and are resistant to erlotinib (**Figure 2B, 2C**). Culturing the two clonally-derived lines in the presence of DOX resulted in a 4 to 8-fold increase of *KMT5C* relative to cells grown in PBS containing media (**Figure 4A**). H4K20me3 levels were also significantly increased following DOX induction in both clones, but not in Calu6 parental cells (**Figure 4B**). Exposure of clones to increasing concentrations of erlotinib resulted in ~2-fold increase in GI50 values for clones cultured in DOX (**Figure 4C**). Live-cell proliferation analysis of clone 2 in the presence of three different concentrations of erlotinib validated these findings (**Figure 4D**). With respect to gefitinib, afatinib and osimertinib, SUV420H2 overexpressing clones were sensitized (**Supplementary Figure 4**), most notably at higher concentrations of each drug. Evaluation of additional chemotherapeutic and targeted agents in the presence of SUV420H2 are required to validate the role of SUV420H2 in sensitizing resistant cells to other anti-cancer agents.

**Figure 4:**
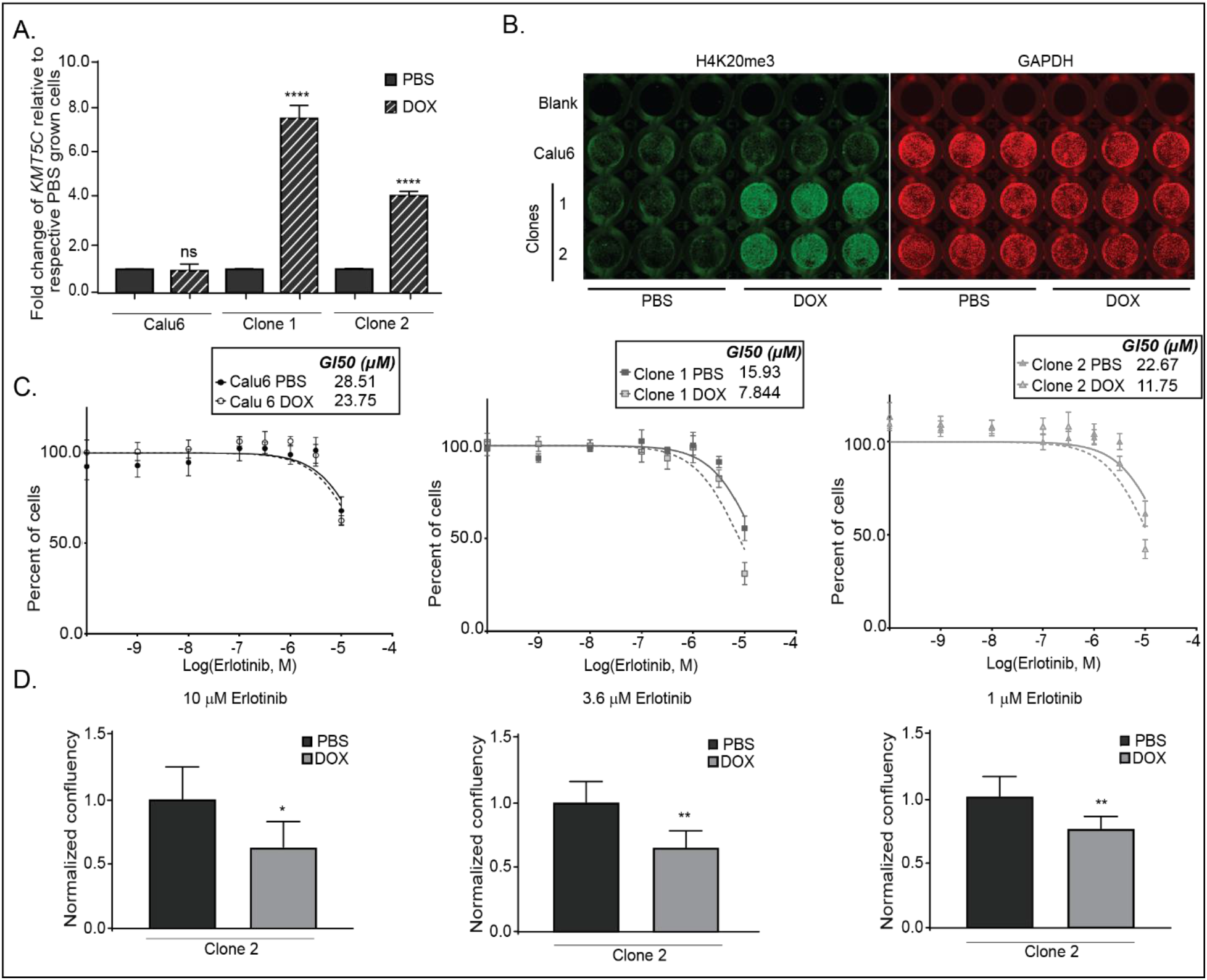
Ectopic expression of SUV420H2 partially sensitizes EGFRi resistant cells to erlotinib. A) SUV420H2 transcript levels evaluated by qRT-PCR in Calu6 cells and Calu6 clones 1, 2 stably expressing DOX-inducible SUV420H2. One-way ANOVA followed by Dunnett’s Multiple Comparison test was used to evaluate statistical significance of SUV420H2 transcript levels relative to respective PBS treated cells. B) H4K20me3 levels evaluated by in-cell western. DOX (or PBS control) treatment was for two weeks. GAPDH serves as an endogenous control. C) Erlotinib dose response measured by SRB was evaluated after a two-week exposure to PBS or DOX containing media. Cells were then exposed to varying concentrations of erlotinib or the highest equivalent volume of DMSO containing media for 72 hours following normalization, the GI50 concentration of erlotinib was calculated from the respective dose curve for each cell line. D) Proliferation of clone 2 was evaluated using the Incucyte. Cells grown in PBS or DOX containing media for two weeks were exposed to varying concentrations of erlotinib or the highest equivalent volume of DMSO containing media for 72 hours. Normalized data relative to respective normalized PBS treated samples is represented. Unpaired t-test was used to evaluate the statistical significance for each pair.

### SUV420H2 negatively regulates the oncogenic long non-coding RNA, LINC01510, and the oncogene, MET

Because SUV420H2 functions as a tumor suppressor, and is associated with repression of oncogenes (Nelson et al., 2016; Shinchi et al., 2015), GEPIA analysis was used to determine if any of the common bypass tracks involved in erlotinib resistance were negatively correlated with *KMT5C*. A significant negative correlation was identified between *MET* and *KMT5C* in LUAD (Spearman r = −0.44, p-value = 1.0e^−37^ **Supplementary Figure 5A**). MET amplification is one of the more common bypass mechanisms that cells use to overcome inhibition of EGFR signaling by erlotinib (Jakobsen et al., 2017; Liao et al., 2019). As expected, *MET* was higher in LUAD relative to normal tissues (**Supplementary Figure 5B**). To determine if the negative correlation between *MET* and *KMT5C* held true in the NSCLC cell lines, SUV420H2 mutant cells were evaluated for MET. Indeed, following loss of SUV420H2 in the mutant cells, MET was increased relative to levels in SUV420H2 wildtype cells (**Figure 5Ai**). Additionally, the *MET* transcript was elevated in the mutant cells, suggesting that SUV420H2 enhanced MET via a transcriptional mechanism (**Figure 5Bi**). Conversely, induction of SUV420H2 in dox-inducible clones resulted in reductions in both MET RNA and protein (**Figure 5Aii, Bii**).

**Figure 5:**
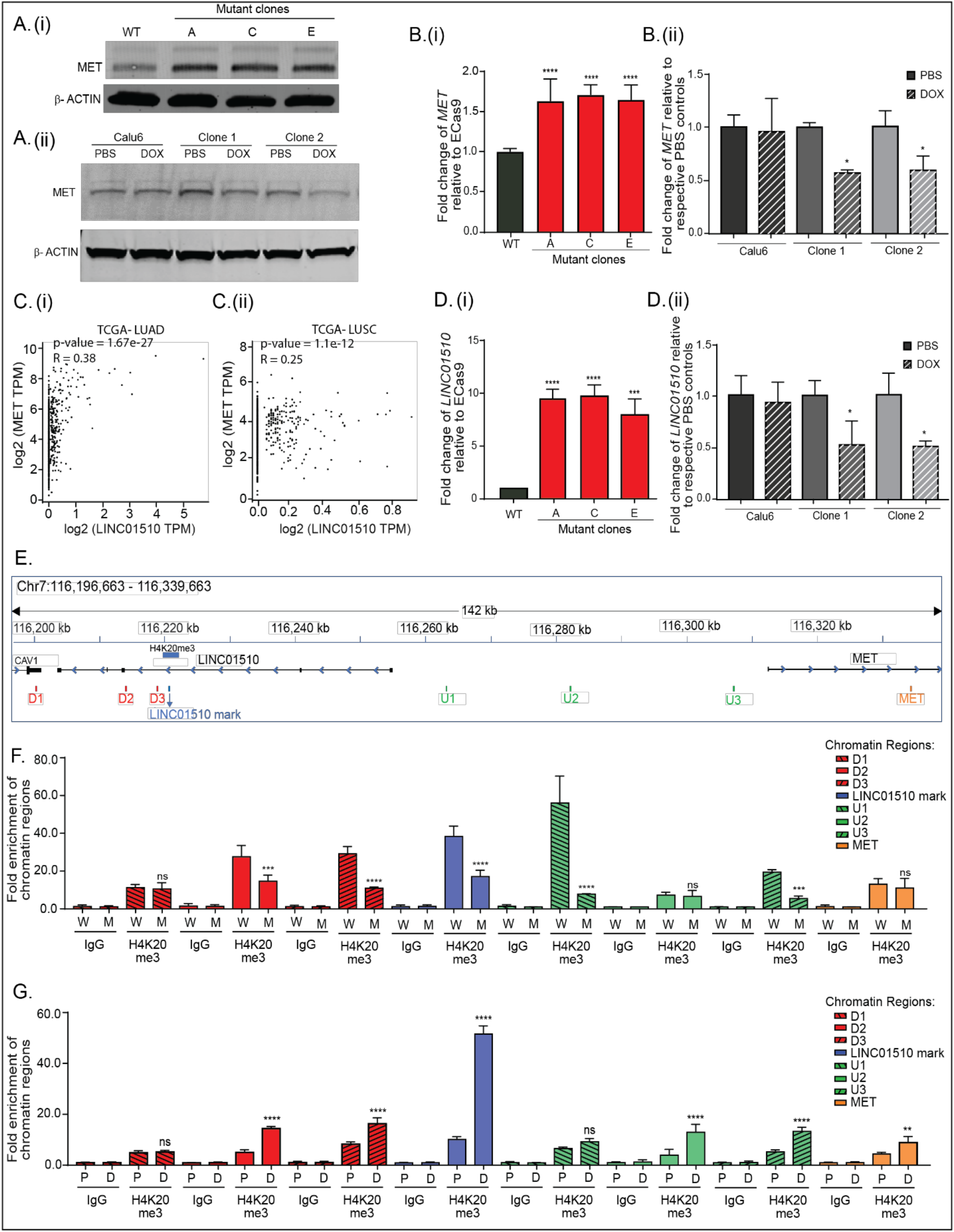
SUV420H2 represses LINC01510 and MET via H4K20me3. A) Representative western blot of MET in (i) SUV420H2 WT cells and mutant clones, (ii) Calu6 cells and clones stably expressing a DOX-inducible SUV420H2 vector. B) qRT-PCR data for *MET* in (i) WT cells and SUV420H2 mutant clones, or (ii) Calu6 cells and clones stably expressing a DOX-inducible SUV420H2 vector. β-ACTIN was used as a loading control for western analysis, qRT-PCR data were normalized to *GAPDH*. C) Correlation analysis between *LINC01510* and *MET* transcript levels obtained from (i) LUAD and (ii) LUSC datasets, evaluated using GEPIA. D) Expression of *LINC01510* in (i) SUV420H2 mutant lines, or in (ii) SUV420H2 inducible clones evaluated by qRT-PCR. E) Diagram of the genomic region (Chr7: 116,196,663 - 116,339,663) representing the predicted H4K20me3 modification on the *LINC01510* gene body, upstream of *MET*, as identified from GSE59316. ChIP-qPCR primers designed on and around the H4K20me3 mark are indicated as LINC01510 mark, regions downstream (D1, D2, D3) and upstream (U1, U2, U3) of the H4K20me3 mark, and on MET. ChIP was performed on chromatin isolated from F) WT (W) or SUV420H2 mutant clone C (M), G) DOX inducible SUV420H2 cells following growth in DOX (D, induced) or PBS (P, uninduced). In F and G either IgG or H4K20me3 primary antibodies were used. qPCR using the immunoprecipitated chromatin was conducted using primers depicted in E (Supplementary Table 3). Data is represented as fold enrichment of the chromatin region pulled-down by H4K20me3 primary antibody relative to IgG. Statistical significance is represented for fold enrichment of chromatin regions in SUV420H2 mutant clone C relative to WT, or DOX relative to PBS. For panels showing statistical significance, One-way ANOVA followed by Dunnett’s Multiple Comparison test was used. TPM= Transcripts per million.

Previous studies determined that MET can be induced through both genomic amplification and transcriptional upregulation (Jakobsen et al., 2017; Pennacchietti et al., 2003; Seol et al., 2000). Although there are multiple mechanisms that are involved in regulating transcription from the *MET* locus, a recent study identified a long non-coding RNA (lncRNA) that functions as a positive regulator of *MET* (Li et al., 2018). A short variant of the long non-coding RNA, LINC01510, referred to as COMET (Correlated-to-MET) was also identified to positively regulate *MET* transcription in papillary thyroid carcinomas (Esposito et al., 2019). Similarly, high LINC01510 correlates with poor prognosis in various cancers, including NSCLC (Li and Wei, 2019; Li et al., 2018, 2019). Further evaluation of *LINC01510* transcript levels in NSCLC via GEPIA analysis in LUAD (TCGA and GTEx) indicated that *LINC01510* was higher in a subset of tumors relative to normal tissues (**Supplementary Figure 5D**).

Since *LINC01510* and *MET* levels are reported to positively correlate in colorectal cancer (Li et al., 2018), their correlation was evaluated in NSCLC. Correlation analyses using TCGA LUAD and LUSC datasets via GEPIA suggested that *LINC01510* and *MET* levels are positively correlated in both LUAD (Spearman r = 0.38, p-value = 1.6e^−27^) and LUSC (Spearman r = 0.25, p-value = 1.1e^−12^) (**Figure 5C**). Based on the reported and evaluated positive correlation between *MET* and *LINC01510*, and the negative correlation between SUV420H2 and MET (**Figure 5 Ai, Bi, C**), we hypothesized that *KMT5C* would also negatively correlate with *LINC01510*. The correlation analysis between *KMT5C* and *LINC01510* suggests a significant, modest negative correlation in LUAD tissues (Spearman r = −0.19, p-value = 1.8e^−7^, **Supplementary Figure 5C).** As also hypothesized, in SUV420H2 mutant clones *LINC01510* was significantly upregulated between 8 and 10-fold (**Figure 5Di**). Conversely, in the SUV420H2 inducible clones, *LINC01510* was significantly lower when cells were cultured in the presence of DOX (**Figure 5Dii**).

SUV420H2 mediates its repressive effects on oncogenes via the H4K20me3 modification (Shinchi et al., 2015), hence we hypothesized that *MET* and/or *LINC01510*, are likely negatively regulated by SUV420H2 via H4K20me3 mediated repression. To this end, we analyzed the reported ChIP-seq profile of H4K20me3 obtained from a human lung fibroblast cell line, IMR90 (GSE59316) (Nelson et al., 2016). Surprisingly the H4K20me3 modification in this dataset was not present within or near the *MET* locus but instead was localized in the gene body of *LINC01510*, i.e. ~55kb upstream of start site of its neighboring gene, *MET* (**Figure 5E**). To identify the region of the chromosome associated with the H4K20me3 modification in the erlotinib sensitive cells, chromatin immunoprecipitation followed by q-RT-PCR (ChIP-qPCR) was conducted. Sensitivity of the assay was first established using primers designed to pulldown the *FOXA1* locus, a target previously reported to be regulated by SUV420H2 via the H4K20me3 modification (Viotti et al., 2018). Two primer sets were tested, one based on the original publication (Viotti et al., 2018), and another that overlaps with the predicted H4K20me3 mark (**Supplementary Figure 6A and Supplementary Table 3**). As expected, pulldown of the FOXA1 region was depended on the presence of SUV420H2. A significant reduction in pulldown was observed in the SUV420H2 mutant clones (**Supplementary Figure 6B**) and an increase in pulldown was evident when SUV420H2 was induced (**Supplementary Figure 6C**).

Following the results obtained from ChIP-qPCR for *FOXA1*, ChIP-qPCR analyses on *LINC01510* and the *MET* loci was conducted using primers that overlapped with the predicted H4K20me3 site and with primers both up and downstream of the predicted site (**Figure 5E**, **Supplementary Table 3**). Similar to the *FOXA1* locus, pulldown varied depending on the status of SUV420H2. The most abundant reduction in pulldown in the SUV420H2 mutant occurred just upstream of the *LINC01510* locus with no obvious difference at the *MET* locus (**Figure 5F** compare upstream primer 1 (U1) to MET primers). In concordance, induction of SUV420H2 followed by ChIP-qPCR resulted in enrichment of the H4K20me3 mark in regions surrounding the lncRNA, with the LINC01510 mark having the most significant increase, with only a marginal increase at the *MET* locus (**Figure 5G**, compare LINC01510 primers to MET primers). Importantly the observed enrichment in ChIP-qRT-PCR for the LINC01510 mark is similar to the enrichment observed at the *FOXA1* locus. These results further support the hypothesis that SUV420H2 regulates *LINC01510* expression via the H4K20me3 modification present on its gene body.

### Loss of LINC01510 or MET partially re-sensitizes SUV420H2 mutant cells to erlotinib, conversely overexpression promotes erlotinib resistance in SUV420H2 wildtype cells

From Figure 5, it can be inferred that SUV420H2 negatively regulates both *LINC01510* and *MET* transcript levels, and MET protein levels. It was also determined that SUV420H2 negatively regulates *LINC01510* transcription, likely via H4K20me3 on its gene body. Therefore, we evaluated if SUV420H2 negatively regulates MET indirectly via repression of *LINC01510*. *LINC01510* or *MET* were knocked down in an SUV420H2 mutant clone, which expresses high levels of *LINC01510* and *MET* (**Figures 5Ai, 5Bi, 5Di**). By western blot and qRT-PCR analyses, it was confirmed that siRNAs targeting either *MET* or *LINC01510* downregulate MET at both the transcript and protein level (**Figure 6A, B**). To determine if loss of SUV420H2 partially mediates erlotinib resistance via upregulation of *LINC01510* and *MET*, *LINC01510* or *MET* were downregulated and erlotinib dose response and proliferation analyses were conducted. Both results validate that erlotinib resistant SUV420H2 mutant cells can be partially re-sensitized to erlotinib post knockdown of either *LINC01510* or *MET* (**Figure 6C, D**). In fact, the more sensitive Incucyte data suggests that re-sensitization is more robust in *LINC01510* knocked down cells than in cells with reduced levels of *MET* (see Figure 6C where *LINC01510* knockdown generates a highly significant reduction in cell proliferation even at low micromolar concentrations of erlotinib).

**Figure 6:**
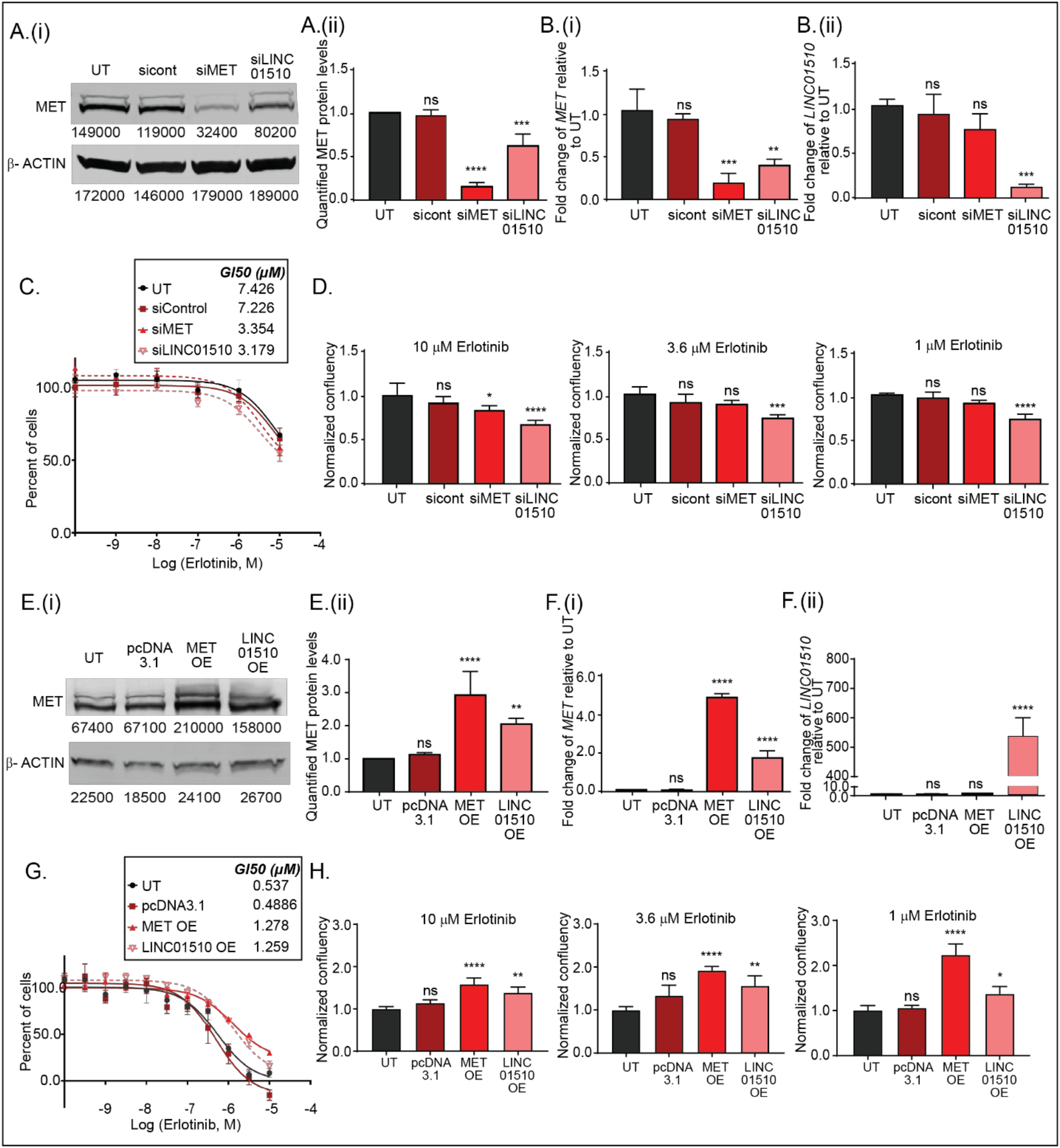
Modulation of LINC01510 or MET are partially responsible for the erlotinib response. A) (i) Representative western blot of MET in SUV420H2 mutant cells that were either untransfected (UT) or reverse transfected with siRNA control (sicont), siRNA to MET (siMET), or siRNA to LINC01510 (siLINC01510) for 96 hours. β-ACTIN serves as a loading control. Densitometry values for the representative blots are shown below. ii) Quantification of protein levels from three biological replicates as done in A. B) Expression of (i) *MET* and (ii) *LINC01510* in SUV420H2 mutant cells that were either UT or reverse transfected with sicont, siMET or siLINC01510 for 96 hours. Data are normalized to GAPDH and are graphed relative to data from UT cells. C) Erlotinib dose response of SUV420H2 mutant cells following transfection with the indicted siRNAs. Twenty-four hours post transfection, cells were exposed to varying concentrations of erlotinib or the highest equivalent volume of DMSO (negative control) containing media for 72 hours. Post-normalization, the GI50 concentration of erlotinib was calculated from each respective dose curve. D) Proliferation of SUV420H2 mutant cells following transfection with the indicted siRNAs. Twenty-four hours post transfection, cells were exposed to varying concentrations of erlotinib or the highest equivalent volume of DMSO containing media for 72 hours. Normalized data is represented relative to that of UT. One-way ANOVA followed by Dunnett’s Multiple Comparison test was used to evaluate statistical significance. E) (i) Representative western blot of MET in SUV420H2 WT cells that were either untransfected (UT) or transfected with pcDNA3.1 control plasmid or plasmids to overexpress to MET (MET OE) or LINC01510 (LINC01510 OE) for 96 hours. β-ACTIN was used as a loading control. Densitometry values for the representative blots are shown below. (ii) Quantification of MET levels from three biological replicates as in A. F) Expression of (i) *MET* and (ii) *LINC01510* in SUV420H2 wildtype (WT) cells that were either UT or transfected with the indicated vectors. Data are normalized to *GAPDH*. G) Erlotinib dose response via SRB assay was evaluated in WT cells that were either UT or that were transfected with the indicated vectors, as described in C. H) Proliferation of WT cells transfected as in G, was evaluated as described in D.

Data presented in Figure 6A, B suggests that knockdown of *LINC01510* reduces MET at the transcript level, therefore, we further evaluated if overexpression of *LINC01510* in SUV420H2 wildtype cells can positively regulate MET. Following transfection of the *LINC01510* plasmid, or *MET* plasmids (positive control), a modest, yet significant increase in MET was observed (**Figure 6E, F**). Additionally, as hypothesized, *LINC01510* or *MET* overexpression also led to acquired resistance in SUV420H2 wildtype cells, evaluated by both dose curve and proliferation analyses (**Figure 6G, H**).

Overall, the findings of this study, depicted in the model in **Figure 7** suggests that wildtype SUV420H2 in NSCLC cells negatively regulates *LINC01510* via the downstream modification, H4K20me3. In cells with high SUV420H2, repression of *LINC01510* inhibits full expression of MET. However, upon loss of SUV420H2 in mutant cells, *LINC01510* becomes de-repressed due to reductions in the H4K20me3 modification, resulting in increased expression of *LINC01510*. Simultaneously, *LINC01510* positively regulates the transcription of *MET*. Therefore, increased levels of *LINC01510* and MET function as mediators of erlotinib resistance in SUV420H2 mutant cells.

**Figure 7:**
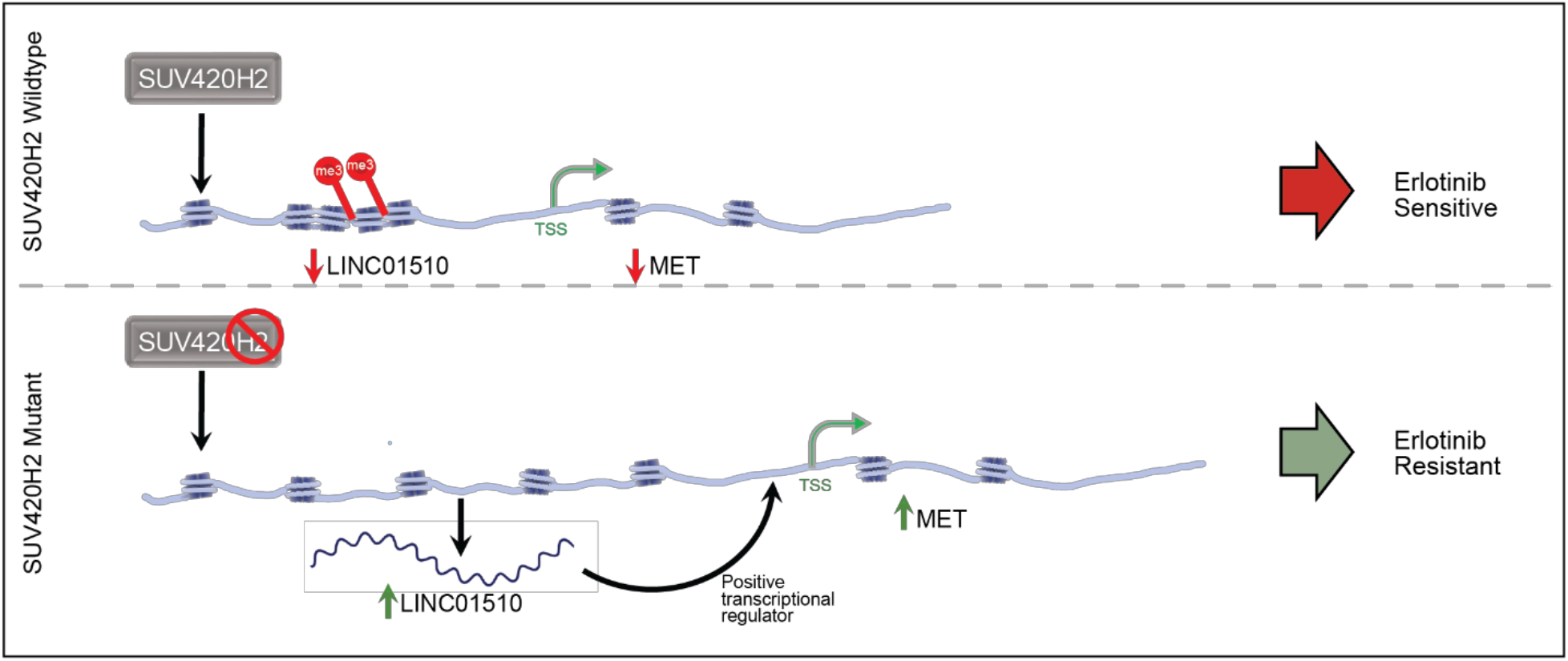
Model depicting loss of SUV420H2 in NSCLC results in development of erlotinib resistance via upregulation of LINC01510 and MET.

## Discussion

Changes to the epigenome are common occurrences that influence all aspects of cancer, including chemoresistance (Flavahan et al., 2017). However, only a limited subset of epigenetic factors have been determined to have a role in resistance to therapeutic drugs in cancer (Wilting and Dannenberg, 2012). The aim of this study was to identify unknown mechanisms by which acquired erlotinib resistance manifests in NSCLC in an unbiased way, and loss of SUV420H2, an epigenetic factor was the top hit. SUV420H2 is a histone methyltransferase responsible for maintaining constitutive heterochromatic regions of the genome and for repressing specific genes, via the repressive mark H4K20me3. Both SUV420H2 and H4K20me3 are significant for maintaining cells in their differentiated states, loss of which is consequentially reported to cause enhanced survival due to elongation of telomeres (Marión et al., 2011), and spontaneous carcinogenesis (Fraga et al., 2005; Pogribny et al., 2007).

Catalysis of H4K20me3 modification of the genome is a sequential process. SUV39H2, another histone methyltransferase first catalyzes the H3K9me3 modification, that further recruits the protein HP1 which physically associates with SUV420H2 to mediate H4K20me3 (Bosch-Presegué et al., 2017; Hahn et al., 2013). Although the findings of this study, for the first time identify a role for SUV420H2 in mediating drug resistance, loss of a key upstream regulator of SUV420H2 activity, SUV39H1/2 has previously been reported to be associated with resistance (Braig et al., 2005; Völkel et al., 2009). SUV39H null mice displayed chromosomal instabilities and increased tumorigenicity (Bosch-Presegué et al., 2017; Hahn et al., 2013; Kovaríková et al., 2018; Peters et al., 2001). Apart from SUV39H1/2, it is also possible that other upstream regulators of SUV420H2 such as HP1 may have an unidentified role in mediating resistance to drugs, such as EGFRi (Bosch-Presegué et al., 2017; Hahn et al., 2013). Indeed, the first identified demethylase for H4K20me3, mineral dust-induced gene (Mdig) was determined to be overexpressed in breast and lung cancer cells antagonizing the effects of the H4K20me3 modification which led to induction of oncogenes (Zhang et al., 2020). Analogous to Mdig overexpression in cancer cells, leading to reduction of H4K20me3, we found that loss of SUV420H2 also leads to depletion of H4K20me3 mark that in turn enhances oncogenes such as LINC01510 and MET.

It has been long appreciated that genomic instability generates tumor heterogeneity and in the presence of a drug, gives rise to resistant cells (Flavahan et al., 2017; Gillies et al., 2012), also a reported mechanism of EGFRi resistance (Nahar et al., 2018; Serizawa et al., 2013). In the current study, complete loss of SUV420H2 function may have led to spontaneous genetic aberrations leading to rapid establishment of resistant population of cells in the presence of erlotinib and other EGFRi. Indeed, previous reports determined that loss of SUV420H1/2 impairs the DDR mechanism, inadvertently leading to accumulation of damaged DNA and increased tumorigenicity (Bromberg et al., 2017; Celeste et al., 2003; Hahn et al., 2013; Jørgensen et al., 2013; Kovaríková et al., 2018; Sanders et al., 2004; Schotta et al., 2008). Therefore, it is possible that in the SUV420H2 mutant cells, where the activity of SUV420H2 is significantly reduced, the chromatin may have suffered *massive* loss of H4K20me3, which disrupted the heterochromatic shield protecting the DNA from damage. On the contrary, in Calu6 cells, which still have modest amounts of H4K20me3 (**Supplementary Figure 2**) the regions of the chromatin lacking H4K20me3 could be localized at oncogenes leading to their upregulation, while the constitutive heterochromatic regions remained marked and compact, preventing genomic instability. Indeed, increased H4K20me3 in Calu6 cells due to DOX-induction of SUV420H2 resulted in reductions in MET and promoted sensitivity to EGFRi (**Figure 4**) suggesting that even modest changes in H4K20me3, or other unidentified mechanisms of SUV420H2 can alter the response of cells to EGFRi. Additional studies addressing the dynamics of SUV420H2 and H4K20me3 and their role in maintaining genomic stability will need to be conducted to support these observations.

While this study defines a role for MET and *LINC01510* upregulation that is mediated by loss of SUV420H2 in EGFRi resistance, there are likely to be several other oncogenes regulated by SUV420H2 that contribute to this phenotype. Using the NCI Cell Miner Database that has sequencing data for the NCI-60 cell lines (Shankavaram et al., 2009), multiple genes previously determined to be involved in NSCLC or in EGFRi resistance were found to be negatively correlated with *KMT5C* (SUV420H2). Some of the top genes negatively correlated with *KMT5C* include Annexin A5 (negative correlation, nc = −0.616), Vimentin (nc = −0.636), *CD44* (nc = −0.637), *AKT3* (nc = −0.612), *PRKD1* (nc = −0.632) a member of the PKC family, *NOTCH* (nc = −0.565), *JUN* (nc = −.0.359) and *ERK* (nc = −0.343) all with p-values <0.01. In this analysis the negative correlation between *MET* and *KMT5C* was −0.337, p-value <0.01. Similar to MET, many of these genes are predicted to contain a H4K20me3 modification as determined using H4K20me3 ChIP from IMR90 (GSE59316) (Nelson et al., 2016) including *AKT, NOTCH, CD44, ERK* and others. It is possible that aberrant SUV420H2 may alter a cohort of genes that could ultimately synergize to promote resistance, similar to the effects observed following aberrant microRNA expression (Adams et al., 2014; Orellana and Kasinski, 2015). Whether or not these additional candidates are also SUV420H2 targets and what their contribution is to resistance remains an active area of investigation.

In conclusion, the results of this study describe that loss of SUV420H2 confers EGFRi resistance in NSCLC cells via a novel mechanism. Loss of SUV420H2 abrogates the H4K20me3 modification at an oncogenic long non-coding RNA, *LINC01510*, resulting in enhanced transcription of *LINC01510*. *LINC01510* in turn functions as a positive transcriptional regulator of the oncogene *MET* consequently resulting in *MET* upregulation, a predominant mechanism of acquired resistance to erlotinib. Therefore, this study establishes a mechanism of erlotinib resistance mediated by loss of SUV420H2, which in part is due to indirect overexpression of *MET*.

## Methods

### Cell culture

All cell lines used in the study were obtained from American Type Culture Collection (ATCC), cultured and were confirmed to be free of mycoplasma. Cell lines generated during the study were authenticated by ATCC Cell Line Authentication. All cell lines were grown in RPMI media supplemented with 10% FBS and 1% Penicillin/Streptomycin. ECas9 cells were continuously cultured in media containing 1μg/ml Blasticidin, SUV420H2 mutant clones were grown in media containing 100ng/ml Puromycin, inducible-SUV420H2 Calu6 clones were cultured in 500ng/ml Puromycin containing media, and rescue clones were grown in media containing 100ng/ml Puromycin and 300μg/ml G418 containing media.

### Drug Preparation for in vitro studies

Erlotinib (S7786, Selleck Chemicals), afatinib (850140-72-6, Sigma Aldrich), gefitinib (S1025, Selleck Chemicals), and osimertinib (S7297, Selleck Chemicals) were dissolved in DMSO to prepare 0.4 M stock solutions, which were aliquoted and stored in −80°C. A 200 μM working dilution of all the drugs was prepared in complete medium and were used to prepare the indicated concentrations for all *in vitro* experiments.

### Knock-out CRISPR screen

EKVX cells (4×10^5^) were plated in 6-well plates and were transfected with 3μg of linearized lentiCas9-Blast (Addgene, 52962) using lipofectamine 2000 (11-668-019, Thermo Fisher Scientific), as per manufacturer’s instructions. Forty-eight hours later, cells were selected using 5μg/ml Blasticidin. ECas9 (clone 7) cells stably expressing Cas9 plasmid were clonally selected and characterized. Lentiviral sgRNA library (A and B) were generated and the titer was determined as previously described (Golden et al., 2017). The GeCKO V2 library has 6 sgRNAs targeting protein coding gene and 4 sgRNAs targeting miRNAs. To achieve a 300-fold coverage of the libraries seventeen 12-well plates were each seeded with 4.5×10^5^ ECas9 cells. Nine plates were transduced with library A, and 8 plates were transduced with library B, both at a multiplicity of infection (MOI) of 0.4 in the presence of Polybrene (10μg/ml). Twenty-four hours post transduction, cells were pooled and ~1.31 × 10^7^ cells were re-plated in each of seven 15 cm plates containing complete media supplemented with 2μg/ml Blasticidin. Forty-eight hours later cells were plated in six 15 cm plates in media containing 2μg/ml Puromycin, to select for library-transduced cells, and 2μg/ml Blasticidin. Seventy-two hours later, 2.6 × 10^7^ cells were stored for baseline and 2.6 × 10^7^ cells were re-plated. The following day, media was replaced with GI75 erlotinib containing media (1.23μM erlotinib) and cells were continuously exposed to GI75 erlotinib for 15 passages. Three biological replicates were performed, and genomic DNA from each baseline and erlotinib treated sample were isolated using the Genomic DNA isolation kit (K1820-01, Thermo Fisher Scientific) following the manufacturer’s protocol. For sequencing library preparation, two sequential PCR reactions were conducted for each sample. The first PCR reaction specifically amplified sgRNAs from 1μg of gDNA isolated from each sample. Twenty-five such PCR reactions were conducted, pooled, and gel purified using QIAEX II Gel Extraction Kit (20021, Qiagen). Each PCR1 reaction product (10 ng) was then used for each of 20 PCR2 reactions that were pooled and gel purified. PCR2 fragment sizes and library quality were evaluated on a bioanalyzer (Agilent). Both PCR1 and PCR2 primers are listed in **Supplementary Table 2** (Integrated DNA Technologies). Barcodes included in PCR2 primers were used to identify the samples after deep sequencing. All sequencing was conducted using a NovaSeq 6000 (Illumina). FastQC version 0.11.7 was used to observe sequencing data quality before and after trimming. Cutadapt version 1.13 was used to trim adapters from reads. Reads post-trimming that were shorter than 18nt were discarded. MAGeCK-VISPR v. 0.5.6 was used to perform mapping, allowing no mismatches to ensure accuracy and to reduce bias. Finally, MAGeCK was used to identify over- and under-represented sgRNAs in treated samples relative to baseline, represented as β-scores (Liu et al., 2014).

### Knockout, knockdown, overexpression and rescue experiments

To generate the SUV420H2 sgRNA, two oligos (see **Supplementary Table 3**) were annealed and 5’ phosphorylated (T4 Polynucleotide Kinase kit, M0201S, NEB) as described previously (LentiGuide-Puro and LentiCRISPRv2). Simultaneously, the CRISPR-Cas9 plasmid, LentiCRISPRv2 (Addgene, 52961) was digested using BsmBI (R0580, NEB), dephosporylated (Antarctic phosphatase, M0289S, NEB) and gel purified using QIAEX II Gel Extraction Kit (20021, Qiagen). The annealed oligos were ligated into the gel purified vector, transformed into Stabl3 bacteria and miniprepped, as outlined previously (LentiGuide-Puro and LentiCRISPRv2). Three micrograms of the so generated pLV-sgSUV420H2 plasmid was linearized and forward transfected in 4×10^5^ ECas9 (SUV420H2 wildtype) cells using lipofectamine 3000 (L3000015, Thermo Fisher Scientific), following the manufacturer’s protocol to generate SUV420H2 mutant clones A, C, E.

For all siRNA-mediated knockdown experiments, 30nM of the respective siRNAs were reverse transfected into 10,000 (for dose curves and proliferation assays) or in 4×10^5^ SUV420H2 mutant clones using Lipofectamine RNAiMAX (13-778-150, Thermo Fisher Scientific) following the manufacturer’s protocol. siRNAs used in the study: siMET (Catalog # 4390824, Assay ID # s8700, Thermo Fisher Scientific) and siLINC01510 (Catalog #: 4392420, Assay ID # n506737 Thermo Fisher Scientific).

For generation of DOX inducible overexpression plasmid, the SUV420H2 sequence was amplified from an ORF expression clone for SUV420H2 (eGFP tagged) (EX-V0810-M98, GeneCopoeia) introducing a stop codon. The sequence was purified and ligated into the pLVX-Tetone. The oligonucleotides used to perform the sequence exchange are indicated in **Supplementary Table 3**. Following construction of the pLVX-Tetone-SUV420H2 plasmid, 3μg of the linearized plasmid was transfected into 4×10^5^ Calu6 cells using lipofectamine 3000 to generate the SUV420H2-inducible Calu6 clone.

Next, to generate the rescue lines from SUV420H2 mutant clone C, a puromycin resistance gene was cloned into pLVX-Tetone-SUV420H2 using the primers outlined in **Supplementary Table 3**. Following generation of the pLVX-Tetone-SUV420H2-puro plasmid, 3μg of the linearized plasmid was transfected in 4×10^5^ SUV420H2 mutant cells using lipofectamine 3000 for the generation of inducible-SUV420H2 rescue clones R1, and R2.

Finally, to test effect of MET or LINC01510 on erlotinib resistance, pT3-EF1a-c-Met (31784, Addgene) or pCMV-Hygro-LINC01510 (Twist Bioscience) were transfected using Lipofectamine 3000 in 4X10^5^ SUV420H2 wildtype cells.

### Genotyping of mutation

Genomic DNA was amplified in the region containing the expected *KMT5C* mutation using Q5 high fidelity polymerase (M0491L, NEB). Primers for amplification and sequencing are outlined in **Supplementary Table 3**.

### Bioinformatic analysis of TCGA data

Cancer Therapeutics Response Portal (CtRPv2) was used to validate the CRISPR-Cas9 knock-out screen (Rees et al., 2016). Gene Expression Profiling Interactive Analysis (GEPIA) database (Tang et al., 2017) (http://gepia.cancer-pku.cn/) was used to evaluate *KMT5C*, *LINC01510*, and *MET* levels in NSCLC patient samples and non-tumorigenic controls. Spearman correlation analysis between *LINC01510* and *MET*, or between *LINC01510* or *MET* and *KMT5C* was also evaluated in LUAD tumor samples using GEPIA. Integrated Genome Viewer (IGV 2.3) was used to view bed files reported by GSE59316 using Human genome 19 (hg19) browser.

### Western Blot

Four-hundred thousand cells were grown in individual wells of a 6 well plate, and lysates were isolated at time points specified in figure legends using RIPA buffer (Sodium chloride (150 mM), Tris-HCl (pH 8.0, 50mM), N P-40 (1 %), Sodium deoxycholate (0.5 %), SDS (0.1 %), ddH2O (up to 100 mL)) containing 1X protease inhibitor cocktail (PIA32955, Thermo Fisher Scientific). Protein quantification was performed using Pierce BCA Protein Assay kit. Equal amounts of protein lysate were resolved through 12% or 4-20% polyacrylamide gels and transferred onto a polyvinylidene difluoride (PVDF) membrane. Membranes were blocked using LI-COR buffer for 1 hour at room temperature, and incubated overnight in primary antibody at 4°C. The primary antibody was detected using 1:800 IR 800CW secondary antibody. Blots were scanned, and data quantified using the Odyssey LI-COR imaging system and software. Antibodies used: mouse H4K20me3 (39672; Active Motif), rabbit H4K20me3 (ab9053, abcam), rabbit MET (D1C2) XP (8198, Cell Signaling), mouse β-ACTIN (3700, Cell Signaling)

### In-Cell Western

Ten-thousand cells were grown in individual wells of a 96-well plate. Forty-eight hours post plating, cells were fixed using cold 100% methanol for 20 minutes at 4 C. Post fixing, cells were permeabilized using 0.2% TritonX in 1X PBS at room temperature for 30 minutes. Cells were blocked using LI-COR blocking buffer for 1.5 hours followed by overnight incubation with primary antibody at 4°C. The primary antibody was detected using 1:800 IR 800CW secondary antibody (LI-COR). The IR-800 signal was quantified using the Odyssey LI-COR imaging system and software. Antibodies used: 1:400 mouse H4K20me3 (39672, Active Motif), 1:500 rabbit GAPDH (2118, Cell Signaling)

### Immunofluorescence

Two-hundred thousand cells were seeded on collagen coated coverslips that were arranged in individual wells of a 12-well plate. Forty-eight hours post plating, cells were fixed using cold 100% methanol for 20 minutes in 4°C. Post fixing, cells were permeabilized using 0.2% TritonX in 1X PBS at room temperature for fifteen minutes. Following which, cells were blocked using LI-COR blocking buffer for 1 hour followed by overnight incubation with 1:50 mouse H4K20me3 (39672, Active Motif) at 4°C. 1:500 anti-mouse Alexa Fluor 647 (A-31571, Thermo Fisher Scientific) was used to detect H4K20me3 and 1:1000 Hoechst dye (H3570, Thermo Fisher Scientific) for 2 hours at room temperature. Coverslips were mounted on glass slides using ProLong Glass Antifade Mountant (P36982, Thermo Fisher Scientific). Fluorescence images were collected using the Nikon A1RMP microscope and analyzed using NIS-Elements Microscope Imaging Software.

### RNA isolation and Quantitative real time PCR (qRT-PCR)

Four-hundred thousand cells were grown in individual wells of a 6-well plate, and total RNA was isolated after 48 or 96 hours, as indicated, using the miRneasy Kit (217004, Qiagen) according to the manufacturer’s instruction. DNase I digestion (79254, Qiagen) was used in each RNA purification reaction to remove genomic DNA. RNA integrity was evaluated on a 1.5% agarose gel, and total RNA quantified using a nanodrop. cDNA was then synthesized from 1μg of total RNA using MiScript Reverse Transcriptase kit (218161, Qiagen), as indicated by the manufacturer’s protocol. Q-RT-PCR was conducted using the miScript SYBR Green PCR Kit (218073, Qiagen) as indicated by the manufacturer’s protocol, to quantify target gene mRNA expression. The following primers were obtained: *GAPDH* (loading control) (QT00079247, Qiagen), *LINC01510* (LPH09040A, Qiagen), and MET (QT00023408, Qiagen). Primers for *KMT5C* quantification are indicated in **Supplementary Table 3**.

### ChIP-qPCR

Briefly, a total of 2×10^7^ cells were fixed using 1% of filter-sterilized formadehyde for 10 minutes at room temperature. The formaldehyde was quenched with 2.5M Glycine (55μL per ml of media) for 5 min. Cells were washed with cold PBS and scraped into fresh cold PBS. Cells were pelleted by centrifuging at 1500 rpm for 10 minutes at 4°C. The cell pellet was resuspended in 10 mL of freshly prepared cold cell lysis buffer (5mM PIPES, 85mM KCl, 0.5% NP40), kept on ice for 10 minutes followed by centrifuging at 1000 rm for 10 minutes at 4°C. The lysed cells were resuspended in 1 mL of nuclei lysis buffer (50mM Tris-HCl (pH 8.0),10mM EDTA, 1% SDS) containing 0.1% protease inhibitor cocktail (PIA32955, Thermo Fisher Scientific) and were transferred into 2mL eppendorf tubes, on ice. Cross-linked chromatin from the isolated nuclei was sonicated using a probe sonicator (60% duty cycle) for 10 seconds with a 1 minute rest, for 15 cycles to fragment DNA (100-500 bps). Fragmented DNA was immunoprecipitated with antibodies against mouse H4K20me3 (39672, Active Motif), or negative control mouse IgG (5415, Cell Signaling Technology) at 4°C overnight with gentle rotation. The immunoprecipitated DNA was purified using the DNA isolation kit (K1820-01; Thermo Fisher Scientific) following manufacturer’s protocol. DNA was used as a template for qRT-PCR as described above. All primer sequences used for qRT-PCR are listed in **Supplementary Table 3**. ChIP data are presented as fold enrichment of DNA immunoprecipitated with H4K20me3 relative to values obtained for DNA immunoprecipitated with IgG control.

### Erlotinib dose response

The protocol followed to evaluate erlotinib dose response was as per the NCI-60 Cell Five-Dose Screen (NCI-60 DTP). Briefly, Sulforhodamine B colorimetric assay (SRB assay)(Orellana and Kasinski, 2016) was performed by exposing cells to varying concentrations of erlotinib or the highest equivalent volume of DMSO (negative control) containing media for 72 hours. To normalize data, percent of cells was calculated based on first correcting for the number of cells at the start of the assay (time zero = tz), followed by normalization of cell number to respective corrected DMSO values.

### Proliferation

Ten thousand NSCLC cells or transfected cells were seeded in 6 replicates in wells of a 96-well plate, which was placed in a live-imaging system, Incucyte s3 2018A (ESSEN BioScience). Plates were incubated in the system for the specified times. Four images per well were obtained every 2 hours using the 10X objective. Confluence was evaluated using Incucyte s3 2018A software. To normalize data, percent of cells was calculated based on first correcting for the number of cells at the start of the assay (time zero = tz), followed by normalization of cell number to respective corrected DMSO values. Data is represented relative to controls, as described in figure legends.

### Statistical analysis

All data were analyzed using GraphPad Prism version 7 software (GraphPad Software) and are presented as mean values ± standard deviation (SD). Pearson’s correlation was utilized to evaluate linear correlation between SUV420H2 and/or H4K20me3 and GI50 erlotinib values. Student’s t-test or one-way ANOVA were performed, as specified in the figure legends. P-value of < 0.05 was considered significant.

## Supplemental Information

Supplementary information can be found online at

## Acknowledgments

The authors gratefully acknowledge the support of multiple core facilities from the Purdue Center for Cancer Research, NIH grant P30CA023168. Dr. Wen-Hung Wang, Genetics Core Facility Director, Bindley BioScience Center, Purdue University for his generous help during SUV420H2 inducible plasmid generation. Dr. Phillip SanMiguel, Genomics Facility Director, Purdue Agriculture, Purdue University for his guidance during deep-sequencing of the CRISPR-Cas9 knock-out screen samples. Dr. Andy Schaber, Imaging Facility Director, Bindley BioScience Center for his inputs during conduction of immunofluorescence experiments. This study was funded in part by the National Institutes of Health (R01CA205420); A.S.P. was supported by a Purdue Research Foundation (PRF) Research Grant award by the Department of Biological Sciences, Purdue University, a SIRG grant administered through the Purdue Center for Cancer Research, Purdue University, a Cancer Prevention Internship Program Graduate Research Assistantship funded by Purdue University, and a Bilsland Dissertation Fellowship awarded by the Department of Biological Sciences, Purdue University.

## Author Contributions

A.S.P. and A.L.K. conceived of the study and developed the methodology. A.S.P. performed the CRISPR-Cas screen. A.S.P and A.M.A. performed validation experiments and analyzed data. N.A.L. performed the bioinformatic analysis. A.S.P and N.A.L. curated the data. A.S.P., A.M.A., J.C., and K.G. validated results/experiments. A.S.P. and A.L.K. wrote and edited the manuscript. A.L.K. supervised the entire project.

