## Supplemental Material for "Loss of SUV420H2 promotes EGFR inhibitor resistance in NSCLC through upregulation of MET via LINC01510"

**Supplemental Information:**

Supplementary Figure 1: Characterization of Cas9 expressing EKVX clones

Supplementary Figure 2: Reduced H4K20me3 correlates with erlotinib resistance in NSCLC cells.

Supplementary Figure 3: SUV420H2 knock-out confers resistance to various EGFRi.

Supplementary Figure 4: Ectopic expression of SUV420H2 partially sensitizes resistant cells to additional EGFRi.

Supplementary Figure 5: LINC01510 correlates poorly with LUAD prognosis.

Supplementary Figure 6: H4K20me3 is enriched at the FOXA1 locus in an SUV420H2 dependent manner.

Supplementary Table 1: Primer sequences used to conduct the CRISPR-Cas9 knock-out screen.

Supplementary Table 2: Primers utilized in the study.

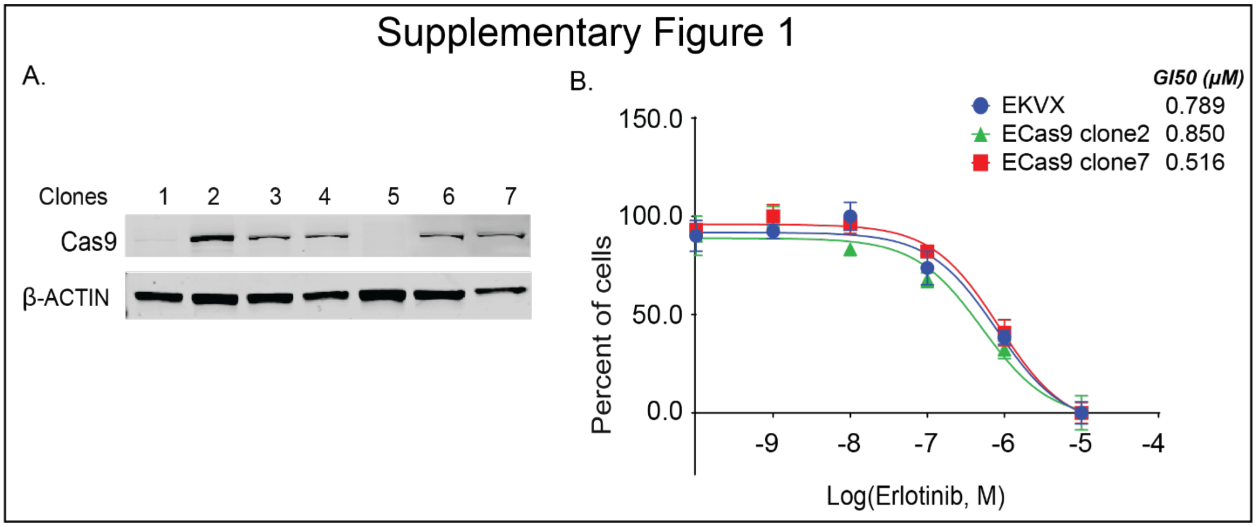


**Supplementary Figure 1: Characterization of Cas9 expressing EKVX clones.** A) Western Blot analysis of Cas9 levels in EKVX clones stably expressing Cas9. β-ACTIN was used as a loading control. B) Parental EKVX cells, ECas9 clone 2, and ECas9 clone 7 were exposed to varying concentrations of erlotinib or the highest equivalent volume of dimethyl sulfoxide (DMSO, negative control) containing media for 72 hours. Erlotinib dose response was evaluated using the SRB assay. Post-normalization, the GI50 concentration of erlotinib was calculated from the respective dose curve for each cell line.

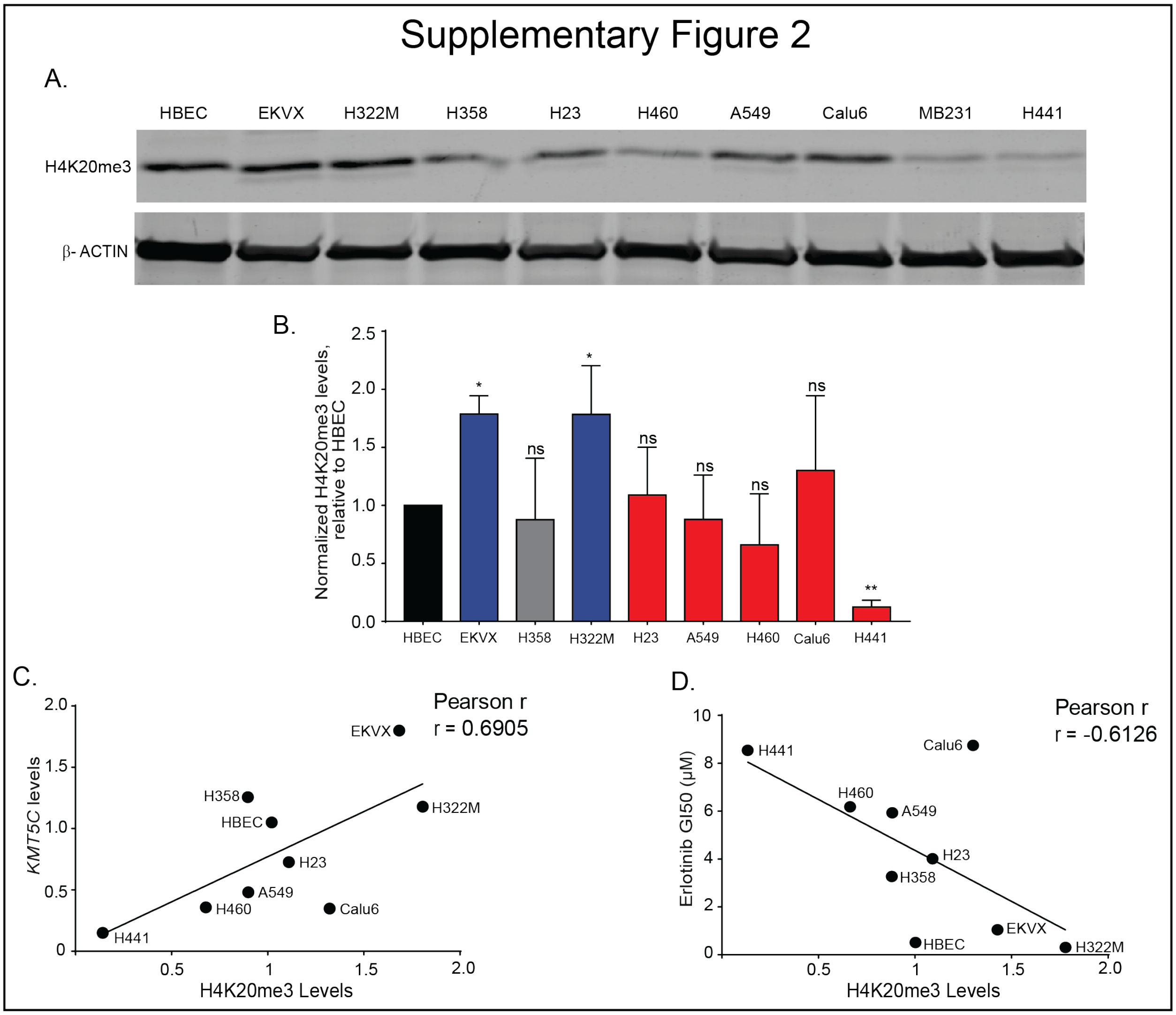


**Supplementary Figure 2: Reduced H4K20me3 correlates with erlotinib resistance in NSCLC cells.** A) Representative western blot of H4K20me3 in a panel of NSCLC cells. β-ACTIN was used as a loading control. MB231, a breast cancer cell line was included as a control cell line reported to have low levels of SUV420H2 (Shinchi et al., 2015). B) Protein quantification of H4K20me3 levels relative to β-ACTIN normalized to HBEC (n=4). One-way ANOVA followed by Dunnett's Multiple Comparison test was used to evaluate statistical significance of H4K20me3 relative to HBEC cells. C) Correlation analysis between quantified H4K20me3 levels from panel B and GI50 erlotinib values from Figure 2B. D) Correlation analysis between quantified H4K20me3 levels from panel B and *KMT5C* from Figure 2A. Evaluations in C and D were conducted using the Pearson correlation test.

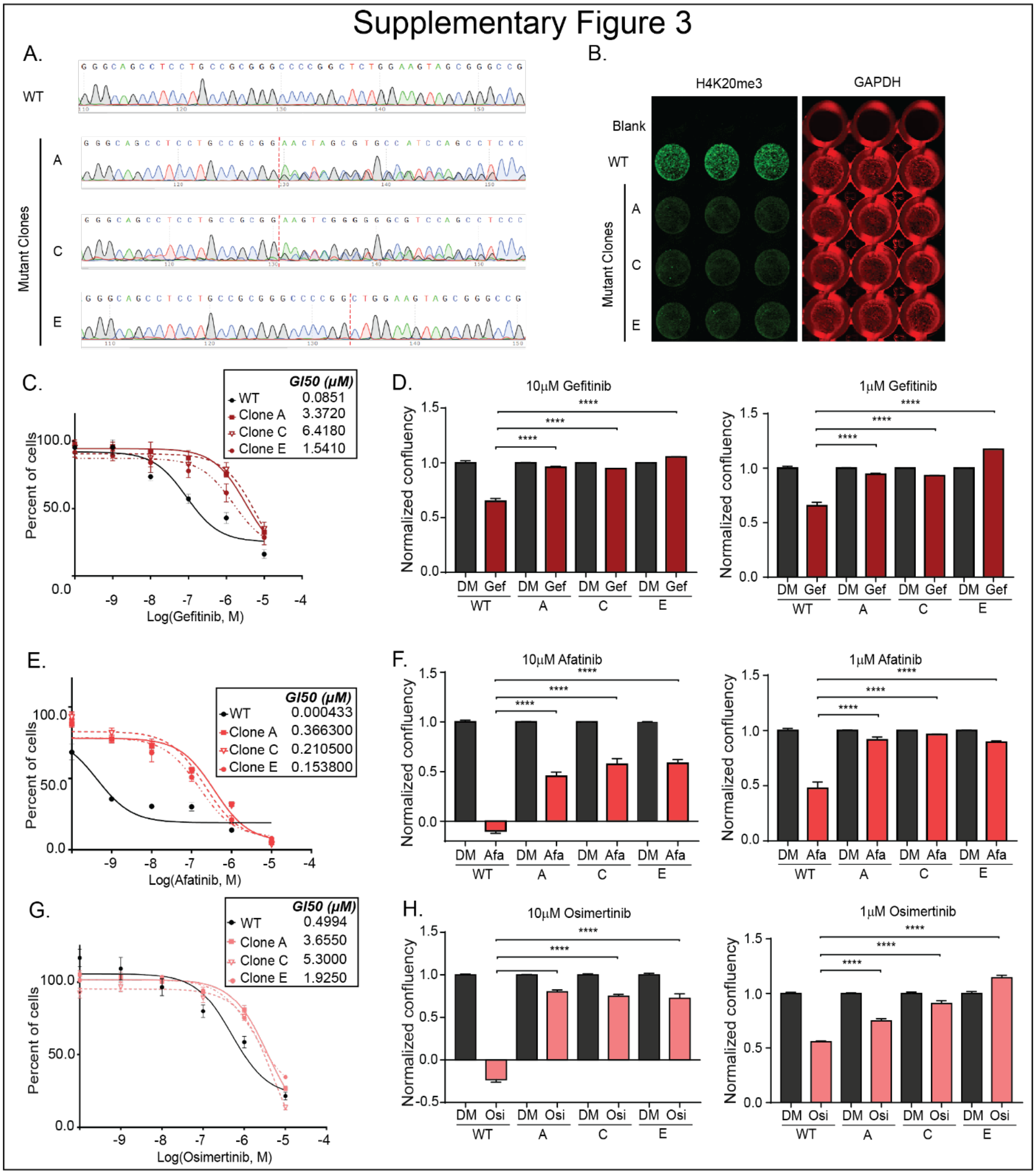


**Supplementary Figure 3: SUV420H2 knock-out confers resistance to various EGFRi.** A) Genomic DNA of WT cells or mutant clones A, C, E was isolated, the region targeted by CRISPR-Cas9 sgRNA targeting SUV420H2 was PCR amplified, purified and sequenced. Representative chromatograms of the wildtype SUV420H2 (WT) cells, and the specific mutations identified in mutant clones A, C, E. B) In-cell western of H4K20me3 levels in WT cells and mutant clones A, C, E. GAPDH serves as an endogenous control. C) Gefitinib, E) Afatinib, or G) Osimertinib dose response curves. Cells were exposed to the indicated concentration of drug or to the highest equivalent volume of vehicle control containing media for 72 hours. Following normalization, the

GI50 concentration of each inhibitor was calculated from the respective dose curve for each cell line. Proliferation of WT cells or mutant clones A, C, E was evaluated using the Incucyte. Cells were exposed to varying concentrations of D) Gefitinib (Gef) F) Afatinib (Afa) or H) Osimertinib (Osi) or the highest equivalent volume of DMSO (DM) containing media for 72 hours. Data relative to respective normalized DMSO control treatments is represented. One-way ANOVA followed by Dunnett's Multiple Comparison test was utilized to evaluate statistical significance of normalized confluency of clones A, C, E in the presence of 10 or 1μM of gefitinib, afatinib or osimertinib compared to WT cells.

Supplementary Figure 4

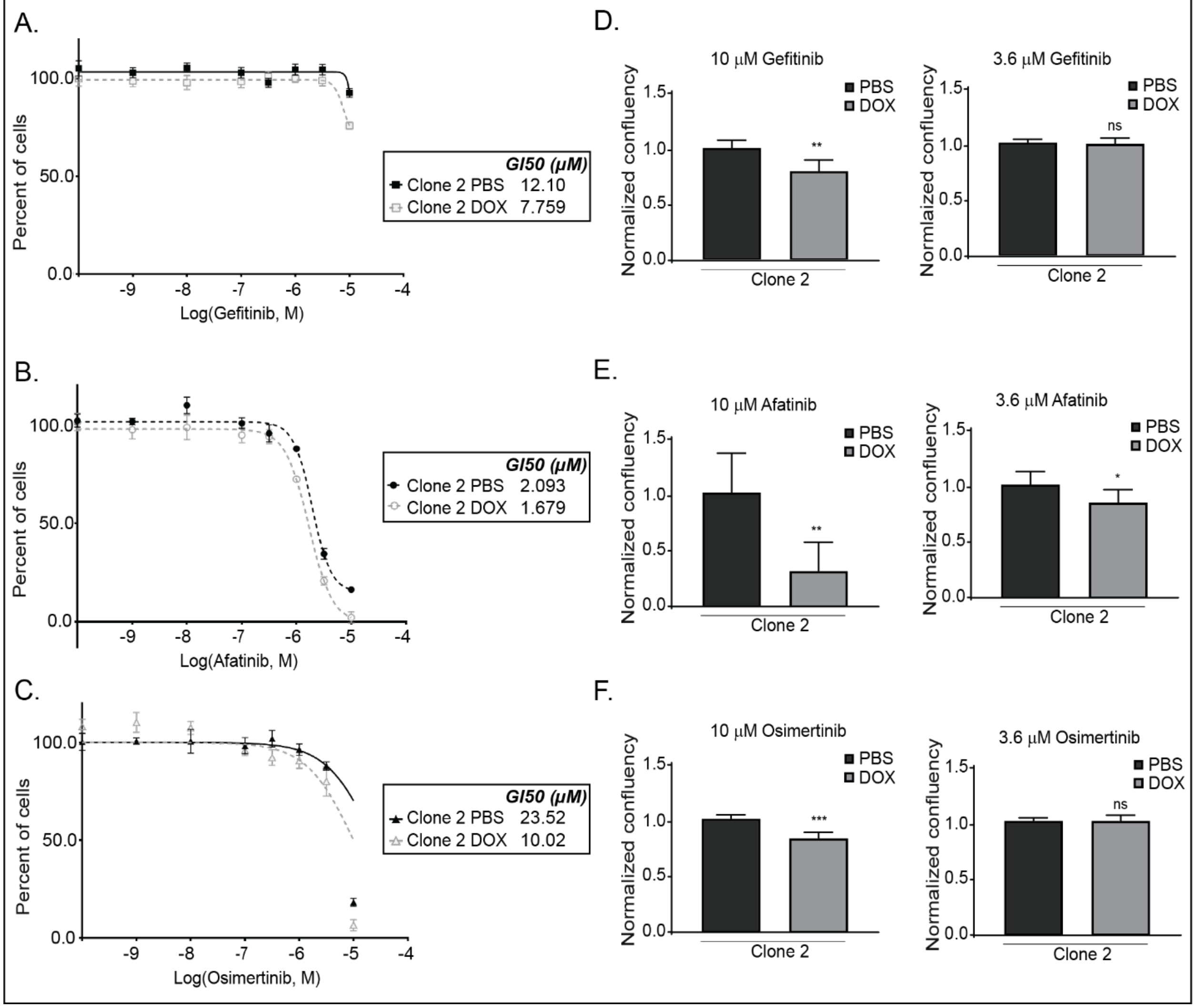


**Supplementary Figure 4: Ectopic expression of SUV420H2 partially sensitizes resistant cells to additional EGFRi.** Dose response measured by SRB was evaluated after a two week exposure to PBS or DOX containing media for Calu6 or clones 1, 2 for A) gefitinib , B) afatininb, C) osimertinib. Erlotinib treatments were for 72 hours. Following normalization, the GI50 concentration of erlotinib was calculated from the respective dose curve for each cell line. Proliferation of clone 2 was evaluated using the Incucyte. Cells grown in PBS or DOX containing media for two weeks, were exposed to varying concentrations of D) gefitinib, E) afatinib, F) osimertinib, or the highest equivalent volume of DMSO containing media for 72 hours. Unpaired t-test was used to evaluate statistical significance of normalized confluency of DOX-cultured clone 2 cells in the presence of either 10 or 3.6 µM of gefitinib, afatinib, or osimertinib compared to respective normalized confluency of PBS-treated cells.

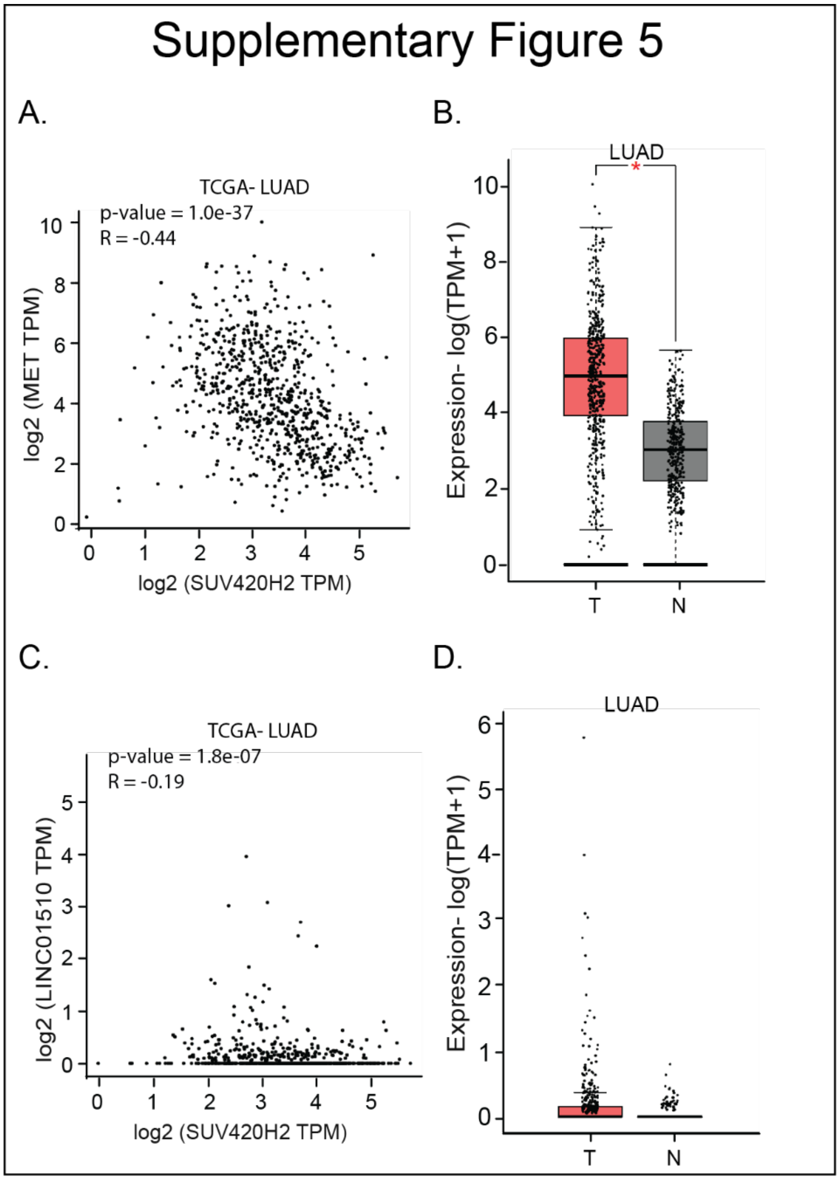


**Supplementary Figure 5: LINC01510 correlates poorly with LUAD prognosis.** Correlation analysis between A) *MET* and *KMT5C* and C) *LINC01510* and *KMT5C* transcript levels in TCGA-LUAD dataset, evaluated using GEPIA.GEPIA analysis for B) *MET* and D) *LINC01510* transcript levels in normal (N, n = 347) and tumor samples (T, n = 483) form LUAD data obtained from TCGA and GTEx databases. The majority of the samples in the normal subgroup had undetectable levels of *LINC01510*. TPM= Transcripts per million

### Supplementary Figure 6

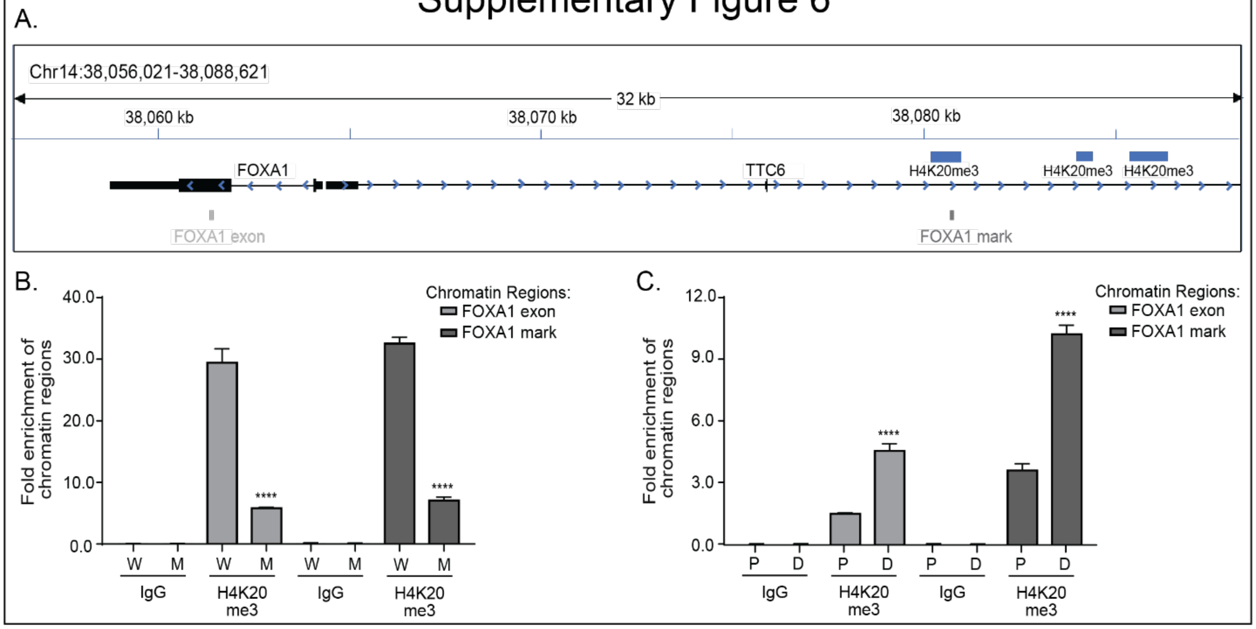


**Supplementary Figure 6: H4K20me3 is enriched at the FOXA1 locus in an SUV420H2 dependent manner.** A) ChIP-qPCR primers designed to evaluate enrichment of H4K20me3 at the FOXA1 exonic region (FOXA1 exon), and at the predicted H4K20me3 modification upstream of the FOXA1 promoter region (FOXA1 mark). ChIP was performed using either IgG or H4K20me3 primary antibodies on chromatin isolated from B) WT or SUV420H2 mutant clone C or C) inducible SUV420H2 cells (in the presence of DOX or PBS). qPCR using the immunoprecipitated chromatin was conducted using primers shown in A (Table 3). Data are represented as fold enrichment of the chromatin region pulled-down by the H4K20me3 primary antibody relative to IgG and was evaluated for significance using one-way ANOVA. W = WT cells, M = SUV420H2 mutant clone C cells, P = Calu6 clones grown in PBS containing media, D = Calu6 clones grown in DOX containing media.

**Supplementary Table 1: Candidate genes identified from the CRISPR-Cas9 knock out screen.** Thirty-five significant hits identified by MAGeCK-VISPR analysis and β-score, p-value, and false discovery rate (FDR)

| **Target** | **β-score** | **p-value** | **FDR** |
|---|---|---|---|
| SUV420H2 | 97 | 8.30E-05 | 0.07 |
| ADSS | 91 | 0.00021 | 0.07 |
| OPA3 | 89 | 0.00028 | 0.07 |
| LEPREL4 | 88 | 0.00032 | 0.07 |
| GAREM | 86 | 0.00049 | 0.07 |
| ISG15 | 83 | 0.00065 | 0.07 |
| PROM2 | 83 | 0.00065 | 0.07 |
| hsa-mir-602 | 77 | 0.00082 | 0.07 |
| CCDC130 | 81 | 0.00088 | 0.07 |
| PCSK2 | 80 | 0.00091 | 0.07 |
| FAM120AOS | 79 | 0.001 | 0.07 |
| CCL23 | 79 | 0.0011 | 0.07 |
| TNFSF12 | 76 | 0.0028 | 0.07 |
| hsa-mir-27b | 74 | 0.0081 | 0.11 |
| SMN2 | 25 | 0.012 | 0.16 |
| OR6V1 | 74 | 0.012 | 0.16 |
| SYBU | 72 | 0.012 | 0.17 |
| CASP8 | 73 | 0.012 | 0.17 |
| LDLRAP1 | 71 | 0.013 | 0.17 |
| PFDN2 | 70 | 0.013 | 0.17 |
| CPA3 | 68 | 0.013 | 0.17 |
| PP2D1 | 68 | 0.013 | 0.17 |
| TMEM234 | 68 | 0.013 | 0.17 |
| TMEM147 | 67 | 0.013 | 0.17 |
| hsa-mir-5699 | 62 | 0.016 | 0.21 |
| hsa-mir-512-1 | 50 | 0.016 | 0.21 |

| | | | |
|---|---|---|---|
| MLL2 | 22 | 0.016 | 0.21 |
| hsa-mir-648 | 43 | 0.016 | 0.21 |
| AGAP9 | 22 | 0.016 | 0.21 |
| hsa-mir-4669 | 43 | 0.016 | 0.21 |
| RPL41 | 38 | 0.016 | 0.21 |
| hsa-mir-3183 | 37 | 0.016 | 0.21 |
| hsa-mir-1268a | 34 | 0.017 | 0.22 |
| hsa-mir-147b | 34 | 0.017 | 0.22 |
| hsa-mir-148a | 27 | 0.018 | 0.24 |

**Supplementary Table 2: Primer sequences used to conduct the CRISPR-Cas9 knock-out screen.** Multiple PCR2 primers were used, each with an independent barcode that allows for sorting of sample-specific sgRNAs post sequencing.

| **PCR** | **Sample** | **Primer name** | **Primer direction** | **Primer sequence** |
|---|---|---|---|---|
| PCR 1 | All samples | 1st PCR primer | Forward | TCTTTCCCTACACGACGCTCTTCCGATCTNNNNAATGGACTATCATATGCTTACCGTAACTTGAAAGTATTTCG |
| | | 1st PCR primer | Reverse | GTGACTGGAGTTCAGACGTGTGCTCTTCCGATCTNNNNGCACCGACTCGGTGCCACTTTTTCAAGTTGATAACGGACTAGCC |
| PCR2 | EKVX-Baseline 1 | UDA5050 | Forward | AATGATACGGCGACCACCGAGATCTACACTGACAATGTCACACTCTTTCCCTACACGAC |
| | | UDA7143 | Reverse | CAAGCAGAAGACGGCATACGAGATAGAAGCCAATGTGACTGGAGTTCAGACGTG |
| | EKVX-Replicate 1 | UDA5051 | Forward | AATGATACGGCGACCACCGAGATCTACACCGACCTAACGACACTCTTTCCCTACACGAC |
| | | UDA7142 | Reverse | CAAGCAGAAGACGGCATACGAGATGACTCACTAAGTGACTGGAGTTCAGACGTG |
| | EKVX-Baseline 2 | UDA5052 | Forward | AATGATACGGCGACCACCGAGATCTACACTAGTTCGGTAACACTCTTTCCCTACACGAC |
| | | UDA7141 | Reverse | CAAGCAGAAGACGGCATACGAGATAGTCTGTCGGGTGACTGGAGTTCAGACGTG |
| | EKVX-Replicate 2 | UDA5053 | Forward | AATGATACGGCGACCACCGAGATCTACACGCCGCACTCTACACTCTTTCCCTACACGAC |

| | | | | |
|---|---|---|---|---|
| | | UDA7140 | Reverse | CAAGCAGAAGACGGCATACGAGATGTATTCTCTAGTGACTGGAGTTCAGACGTG |
| | EKVX-Baseline 3 | UDA5054 | Forward | AATGATACGGCGACCACCGAGATCTACACATTATGTCTCACACTCTTTCCCTACACGAC |
| | | UDA7139 | Reverse | CAAGCAGAAGACGGCATACGAGATACGCCTCTCGGTGACTGGAGTTCAGACGTG |
| | EKVX-Replicate 3 | UDA5055 | Forward | AATGATACGGCGACCACCGAGATCTACACAGAACCGAGTACACTCTTTCCCTACACGAC |
| | | UDA7138 | Reverse | CAAGCAGAAGACGGCATACGAGATTAACCGCCGAGTGACTGGAGTTCAGACGTG |

**Supplementary Table 3: Primers utilized in the study.** Designed and purchased from Integrated DNA Technologies.

| Primer use | | Primer direction | Primer sequence |
|---|---|---|---|
| pLV-sgSUV420H2 | | Forward | CACCGCGGCCCGCTACTTCCAGAGC |
| | | Reverse | AAACGCTCTGGAAGTAGCGGGCCGC |
| pLVX-Tetone-SUV420H2 | | Forward | TCGTAAAGAATTCACCATGGGGCCCGACAGAGTGACAGCA |
| | | Reverse | GAGATCTGGATCCTCAGTACAGCTCTTCACCGCCGAC |
| pLVX-Tetone-SUV420H2-puro | | Forward | CCGCTACGCGTTCAGAAGAACT |
| | | Reverse | AGCGGCGTACGATGATTGAACA |
| *KMT5C* genomic locus amplification | | Forward | GAGCAGATGGGAGGTGCGGCGACAGT |
| | | Reverse | GAGCTCAGAAGAAAGGAGACAGAT |
| *KMT5C* genomic locus sequencing | | Forward | CCTCTCCTTAGCCTGGTCCT |
| | | Reverse | CAAGGGCTAGGAAGTCAGGG |
| *KMT5C* quantification | | Forward | TCGGTTTCCGCACCCATAAG |
| | | Reverse | CGGAGGTAGCGATAGACGTG |
| ChIP-QPCR | FOXA1 mark | Forward | AAGGAGAGGTGCGTTGTTTG |
| | | Reverse | CATTCTCCCACGAAAGGCAG |
| | FOXA1 exon | Forward | AAGACTCCAGCCTCCTCAAC |
| | | Reverse | CGGGTGGTTGAAGGAGTAGT |
| | Linc01510 mark | Forward | GCTTCTTGTCCCTCCCAGAT |
| | | Reverse | GCAGAAGTGAGAGGAAGGGT |
| | Up 1 | Forward | CACACTGGAGTTCTTGCCAC |
| | | Reverse | TATGCACTCCTTCACTGGGG |
| | Up 2 | Forward | GCAGTCCAGCTAAGCAATCC |
| | | Reverse | GACATCTTGGGAAGGGGACA |
| | Up 3 | Forward | CCTCTTCACATCCCACAGGT |

| | | | |
|---|---|---|---|
| | | Reverse | CTCTGCTGGCTTGATCATTG |
| | MET | Forward | GATCAAGGAAATGGGGCGTT |
| | | Reverse | GGGACTAGGGCCTATTGTCA |
| | Down 1 | Forward | CCCTGCCTCTCATCAACTGA |
| | | Reverse | GTTGAGCCACTAAACCACCC |
| | Down 2 | Forward | TGCCTGGTCTCCTGTTAACA |
| | | Reverse | ATCTGTCTTCTCCCTGTGCC |
| | Down 3 | Forward | AGTCCAAGATCAAGGCACCA |
| | | Reverse | AGGCCTTTCTTGTACCCCTT |
